# Quantifying shifts in natural selection on codon usage between protein regions: A population genetics approach

**DOI:** 10.1101/2020.12.18.423529

**Authors:** Alexander L. Cope, Michael A. Gilchrist

**Author notes:** Current: Department of Genetics, Rutgers University Last compiled on Wednesday 8^th^ September, 2021 at 16:24 (Alexander L. Cope).

## Abstract

**Background:** Codon usage bias (CUB), the non-uniform usage of synonymous codons, occurs across all domains of life. Adaptive CUB is hypothesized to result from selection for efficient ribosome elongation, accurate translation, mRNA secondary structure, and/or protein folding. Given the critical link between protein folding and protein function, numerous studies have analyzed the relationship between codon usage and protein structure. The results from these studies have often been contradictory, likely reflecting the differing methods used for measuring codon usage and the failure to appropriately control for confounding factors, such as differences in amino acid usage between protein structures and changes in the frequency of different structures with gene expression.

**Results:** Here we take an explicit population genetics approach to quantify codon-specific shifts in natural selection related to protein structure. As a result, our approach avoids the problem of needing to choose a specific metric of codon usage, naturally takes amino acid usage of a region into account, and explicitly takes variation in gene expression into account. Our results reveal a weak relationship between codon usage and protein structure, indicating that differences in selection between structures are subtle and/or intermittent. While the magnitude of differences in selection are slight, our results indicate the relationship between codon usage and protein structure is more complex than previously believed.

**Conclusions:** While our results clearly indicate that selection on codon usage shifts between secondary structure, the overall impact of these shifts on codon usage are small. Nevertheless, our work demonstrates the statistical power and benefits of studying selective shifts on codon usage or other genomic features from an explicitly evolutionary approach. Limitations of this approach are discussed.

## Background

Patterns of codon usage bias (CUB), or the non-uniform usage of synonymous codons, vary both within and across species [1, 2, 3]. Although non-adaptive evolutionary forces (e.g. mutation biases, GC-biased gene conversion) are well-known to shape codon usage patterns, natural selection also plays a significant role. The correlation between codon frequency and tRNA abundances and the bias towards more efficient codons in highly expressed genes suggests selection against translation inefficiency is a major factor shaping genome-wide codon patterns [4, 5, 6]. Codon usage is also known to vary within a gene, which is hypothesized to reflect various other forces of selection [7]. For example, CUB is thought to be shaped by selection for translation accuracy, such as to reduce missense errors at functionally-important sites and the frequency of ribosomal drop-off later in transcripts, both of which can result in non-functional proteins [8, 9, 10, 11]. Intragenic variation in synonymous codon usage has also been proposed to be shaped by selection to prevent ribosomal queuing [12] and selection to avoid mRNA secondary structure near the ends of mRNA transcripts [13, 14, 15].

Although adaptive CUB is thought to be largely driven by selection for translation efficiency, research indicates potential selective advantages of inefficient codons (“non-optimal” or “rare” codons) at certain sites within a protein [16]. Given that codon usage patterns are strongly shaped by amino acid biases, mutation biases, and gene expression, it is important for researchers investigating possible adaptive codon usage patterns to ensure that these patterns cannot be explained by non-adaptive factors. [17] highlighted the bias of biologists towards adaptationist storytelling, arguing that non-adaptive evolutionary forces (e.g. genetic drift) and other constraints (e.g. developmental) should be considered before attempting to explain a trait or behavior through the lens of adaptive evolution. With the ushering in of the genomic-era, similar arguments have been made about the importance of testing hypotheses related to selection on and adaptation of genomic features in the context of the evolutionary null, i.e. the expectation in the absence of selection [18, 19]. Here, we will use a population genetics based model of coding sequence evolution with clearly defined null formulations to investigate variation in selection on codon usage related to protein structure.

It is generally accepted that misfolded proteins have impaired function, can aggregate within the cell, and possibly disrupt key cellular processes [10, 20, 21]. Missense errors are hypothesized to increase the frequency of protein misfolding; thus, regions important for the protein folding are expected to be under stronger selection for translationally-accurate codons [10]. In addition to the effects of missense errors, codon usage can modulate protein folding via changes in the elongation rates at key steps during cotranslational folding [22]. Empirical evidence indicate changes to elongation rates via synonymous codon usage can alter cotranslational protein folding in organisms ranging from bacteria to multicellular eukaryotes [23, 24, 25, 26, 27, 28]. Synonymous changes to codon usage are known to impact cellular fitness and have been implicated in human diseases through altered protein folding [29, 30]. Understanding the role of codon usage in protein folding also has biotechnological significance, as a recombinant protein is often expressed in an organism with a drastically different CUB, potentially perturbing cotranslational folding [23].

Given that protein secondary structures generally differ in physicochemical properties and ability to cotranslationally fold, researchers have hypothesized that different structures will exhibit different codon usage patterns [16, 31, 32]. Numerous studies have examined the relationship between codon usage and protein secondary structure, but a general relationship, if any, remains unclear [33, 34, 31, 32, 35, 36, 37]. Other analyses of codon usage patterns at higher levels of protein structure often reach different conclusions about the relationship between codon usage and protein structure [38, 39, 37, 40, 32].

Two clear causes for the inconsistencies across studies are (1) the different approaches for quantifying CUB across and within genes and (2) the different ways in which a gene or region is defined as being under selection at the codon usage level. Metrics based on tRNA abundances, such as the tRNA Adaptation Index (tAI) [41], often do not consider relative differences between synonymous codons, but absolute differences across all codons. This makes such metrics particularly prone to amino acid biases when comparing codon usage patterns [16, 42].

Furthermore, while many studies often delineate codons into subsets of “optimal” and “non-optimal” codons, the criterion for classification varies between studies [31, 43]. Indeed, determining codon optimality using tRNA-based metrics has led to the odd situation where all synonyms for an amino acid fall into one group [31]. As we have shown previously [6, 44, 45, 42], even in cases where the codon usage metrics are relative to synonyms for the same amino acid, identifying the optimal codon for an amino acid can be problematic if the effects of mutation and amino acid biases and/or gene expression are not accounted for. Although other studies have attempted to control for these confounding factors, these were often somewhat *ad-hoc* [31].

Recent work has relied on comparative approaches to examine the functional relationship between codon usage and protein structure, recognizing that purifying selection would lead to conserved codon usage patterns [37, 31, 40], although much of this work does not explicitly model evolutionary processes (selection, mutation, drift, etc.) Alternative to species-based comparative approaches are single-genome population genetics approaches which explicitly attempt to model such evolutionary processes. Single-genome population genetics based approaches have been used in various context to examine selection on codon usage [4, 11, 6, 44]. One particularly powerful population genetics approach is the <u>R</u>ibosomal <u>O</u>verhead <u>C</u>ost version of <u>S</u>tochastic <u>E</u>volutionary <u>M</u>odel of <u>P</u>rotein <u>P</u>roduction <u>R</u>ates (ROC-SEMPPR), which is able to separate out the effects of mutation and selection on codon usage by accounting for the natural variation in intergenic gene expression [6, 44, 45]. Unlike many approaches which either average codon usage over regions using heuristic metrics or delineate codons categorically as either optimal or non-optimal, ROC-SEMPPR provides quantitative, codon-specific estimates of mutation bias and natural selection. More specifically, the estimates of selection for each codon from ROC-SEMPPR reflect the population genetics parameter *sN_e_* – the selection coefficient of a codon times the effective population size – in a gene of average expression.

ROC-SEMPPR was originally developed for estimating selection and mutation biases based on genome-wide codon frequencies, but recent work has used ROC-SEMPPR to investigate both intragenic and intragenomic differences in codon usage patterns [42, 46]. ROC-SEMPPR is implemented in a Bayesian framework [44, 47], allowing for model comparisons using Deviance Information or similar criteria. As a proof of principle, we tested for differences in selection on codon usage related to protein secondary structures and intrinsically-disordered regions (IDRs) in *S. cerevisiae* and *E. coli*, two common model organisms for studying CUB. Although model comparisons indicate selection on codon usage differs across protein structures in both species, these differences are relatively small quantitative differences rather than general or qualitative shifts in the nature of selection on codon usage between protein structures. In other words, for both *S. cerevisiae* and *E. coli*, natural selection on codon usage across protein structures is largely consistent, with differences in selection related to folding of different categories of protein structures likely being rare, weak, or both. This highlights a key point that was sometimes missing from previous analyses: although differences in codon usage across protein structures may be statistically significant and even reflect selective differences (assuming the proper controls are used), these effects are overall very small. Based on our results, claims that certain structures “preferentially” use “non-optimal”, “rare”, or “slow” codons are overstated [36, 39]. Quantitative shifts in selection are more consistent with claims that non-optimal codons are “enriched” in some protein structures relative to others (again, assuming the proper controls are used).

Similar to the slight shifts in selection between secondary structures, we also find evidence for slight position-specific shifts in natural selection within helical regions in both *S. cerevisiae* and *E. coli*. However, we show that a previously detected enrichment of slow translating codons near the start of helices, which was proposed to be due to selection to assist in the cotranslational folding [31], was the result of biases in amino acid usage and/or failing to control for the effects of gene expression. Overall, this work demonstrates the power of population genetics approaches for testing hypotheses related to intragenic differences in selection on codon usage.

## Results

### Test of method

Previous work has made claims regarding qualitative differences in the nature of selection on codon usage related to protein structure [36, 39]. Using a simulated dataset based on the empirically-determined protein secondary structures from *S. cerevisiae* (1,097 genes), we tested for qualitative differences in natural selection between protein regions (see Methods for details). Insight into ROC-SEMPPR’s behavior can be gained by observing the effects of having half of the codons evolving under the opposite selective pressure from the remaining codons In this case, ROC-SEMPPR is unable to identify the selectively-favored codon in Region B, leading to a flat line when comparing selection estimates Δ*η* between Region A and Region B. The observed differences in simulated data indicate the estimates of natural selection Δ*η* essentially reflect a weighted average of the various selective forces shaping coding sequence evolution within a region. As noted elsewhere, the Δ*η* value of a codon is equal to *sN_e_* value for that codon relative to the most selectively favored codon of an amino acid when encoded in a gene with average expression, i.e. *φ* = 1. We were able to detect overall selective differences between two regions, even if only 1% of sites in one of the regions was shaped by a different selective force, e.g. selection for inefficiency instead of against inefficiency (Additional File 1, S1). Our simulated results likely represent an approximate lower bound on the number of sites under differing selective pressures necessary to detect <u>qualitative</u> differences in natural selection between protein structures using ROC-SEMPPR. Furthermore, simulations also indicate our approach for detecting codon-specific quantitative shifts has a low false positive rate (Additional File 1, Figure S2).

### Selection on codon usage varies between protein secondary structures

Based on predicted secondary structures from PsiPred [48] in *S. cerevisiae*, we found that the best supported model allowed selection on codon usage to differ across helices, sheets, and coils (Table 1, Model Y_1_). This model is 68 Deviance Information Criterion (DIC, see Methods for details) units better than the next best model assuming no difference in selection between helices and sheets (Y_2_), and 621 DIC units better than the null model assuming no difference across secondary structures. We obtained similar results when using empirically-determined secondary structures, but the best two models were ambiguous as to whether selection differed between helices and sheets (Additional File 1, Table S1 Y_1_ vs Y_2_, ΔDIC *≈* 1).

**Table 1.** Comparison of model fi examining variation in codon usage between predicted pro- tein secondary structures. The null model (Y0) assumes no diff in selection between secondary structures). H: helix. E: sheet. C: coil. ΔDIC = DIC_i_ − DIC_Best_.

| Species | Model | Groupings | | | $\Delta\text{DIC}$ |
| --- | --- | --- | --- | --- | --- |
|  |  | I | II | III |  |
| *S. cerevisiae* | $Y_1$ | H | E | C | 0 |
| | $Y_2$ | HE | – | C | 68 |
| | $Y_3$ | H | EC | – | 285 |
| | $Y_4$ | HC | E | – | 425 |
| | $Y_0$ | HEC | – | – | 621 |
| *E. coli* | $E_1$ | H | E | C | 0 |
| | $E_2$ | HE | – | C | 61 |
| | $E_3$ | H | EC | – | 251 |
| | $E_4$ | HC | E | – | 358 |
| | $E_0$ | HEC | – | – | 471 |

Similar results were obtained for *E. coli*. The best overall model fit allowed for selection to differ across helices, sheets, and coils, with a 61 DIC unit improvement over the model assuming no differences between helices and sheets (E_2_) and 471 DIC unit improvement over the null model (E_0_). Unlike *S. cerevisiae*, similar model fits in *E. coli* using empirical data clearly differentiated between the best two models (Additional File 1, Table S2, E_1_ vs E_2_, ΔDIC = 20).

**Table 2:**
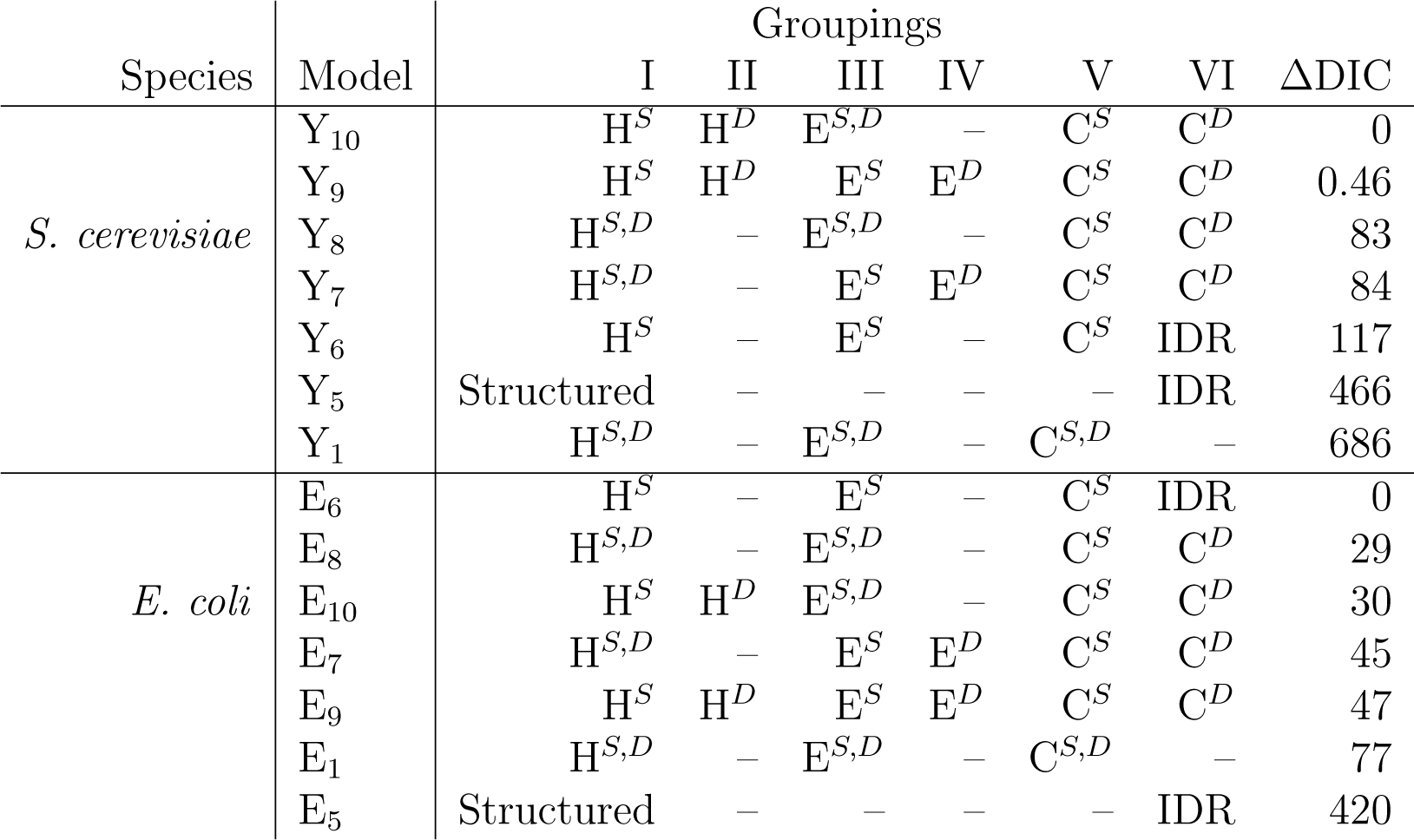
Model comparisons of structure categorizations based on the Deviance Information Criterion (DIC), where the smallest value is considered the best model fi For simplicity, only models which are an improvement over the model over the best secondary structure model (Table 1) are shown, with the exception of the Structured and IDR model for *E. coli*. H: helix. E: sheet. C: coil. Superscripts *S* and *D* indicate if the secondary structure predictions include predictions from structured regions or IDRs, respectively. Structured = H*^S^* E*^S^* C*^S^* . IDR = H*^D^* E*^D^* C*^D^*

### **Comparing selection** Δ*η* on codon usage between secondary structures

Model fits indicate selective shifts on codon usage across protein secondary structures. Comparing selection estimates Δ*η* (based on predicted secondary structures) between protein secondary structures with a Deming regression revealed no significant differences between any of the three secondary structures (Figure 1) in either species, indicating no qualitative selective shifts between any of the three secondary structures.

**Figure 1:**
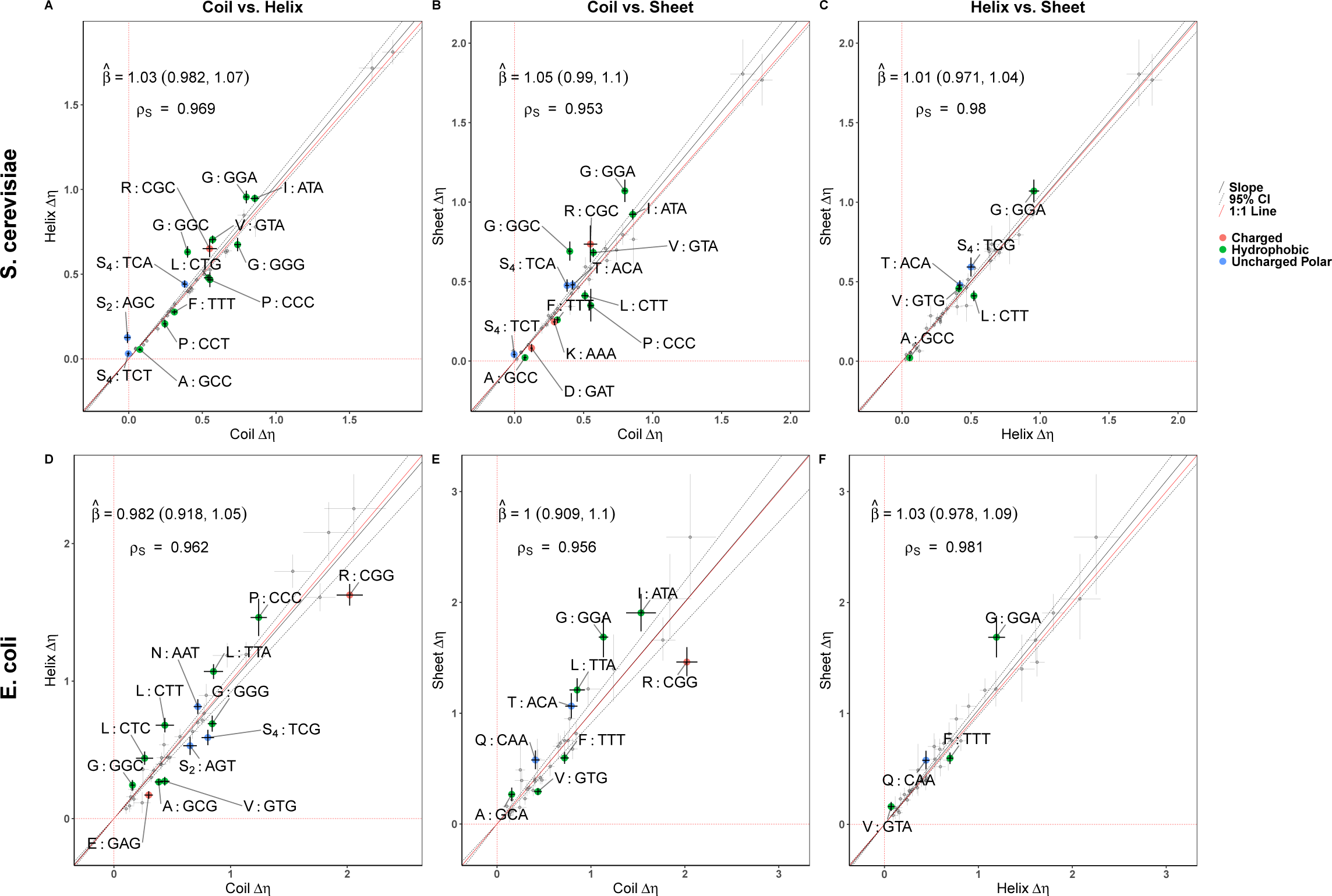
Comparison of selection estimates between diff rent protein secondary structures. values are scaled relative to the genome-wide optimal codon. Points represent values for each codon, while error bars represent the 95% posterior probability intervals. Codons showing significant selective shifts are colored by amino acid property. Negative values indicate a change in the selectively-favored codon relative to the genome-wide optimal. The Deming regression slope and 95% confidence intervals are represented by solid and dashed black lines, respectively. **(A,D)** Coils vs. Helices. **(B,E)** Coils vs. Sheets. **(D,F)** Helices vs. Sheets.

Although no qualitative (i.e. overall) selective shifts on codon usage were detected, examination of the 95% posterior probability intervals for selection estimates Δ*η* indicate clear quantitative differences in the strength of selection related to individual codons. We find that for most of the 18 amino acids with multiple synonyms, selection differs between secondary structures for at least one codon (i.e. its Δ*η* 95% posterior probability intervals do not overlap). These differences mostly reflect quantitative changes in the average strength of selection and not a qualitative switch in the optimal codon. The one qualitative exception to this appears to be serine (S_4_ and S_2_) in coils of *S. cerevisiae*. While codons TCT and AGC are disfavored in helices and/or sheets, this is not the case for coils in which there appears to be no differences in the preference for these two codons and the genome-wide optimal codons (TCC and AGT, respectively, Figure 1A,B). However, while we do detect quantitative selective shifts across secondary structures, these shifts are very small and are expected to have little impact on codon frequencies across protein secondary structures (Figure 2).

**Figure 2:**
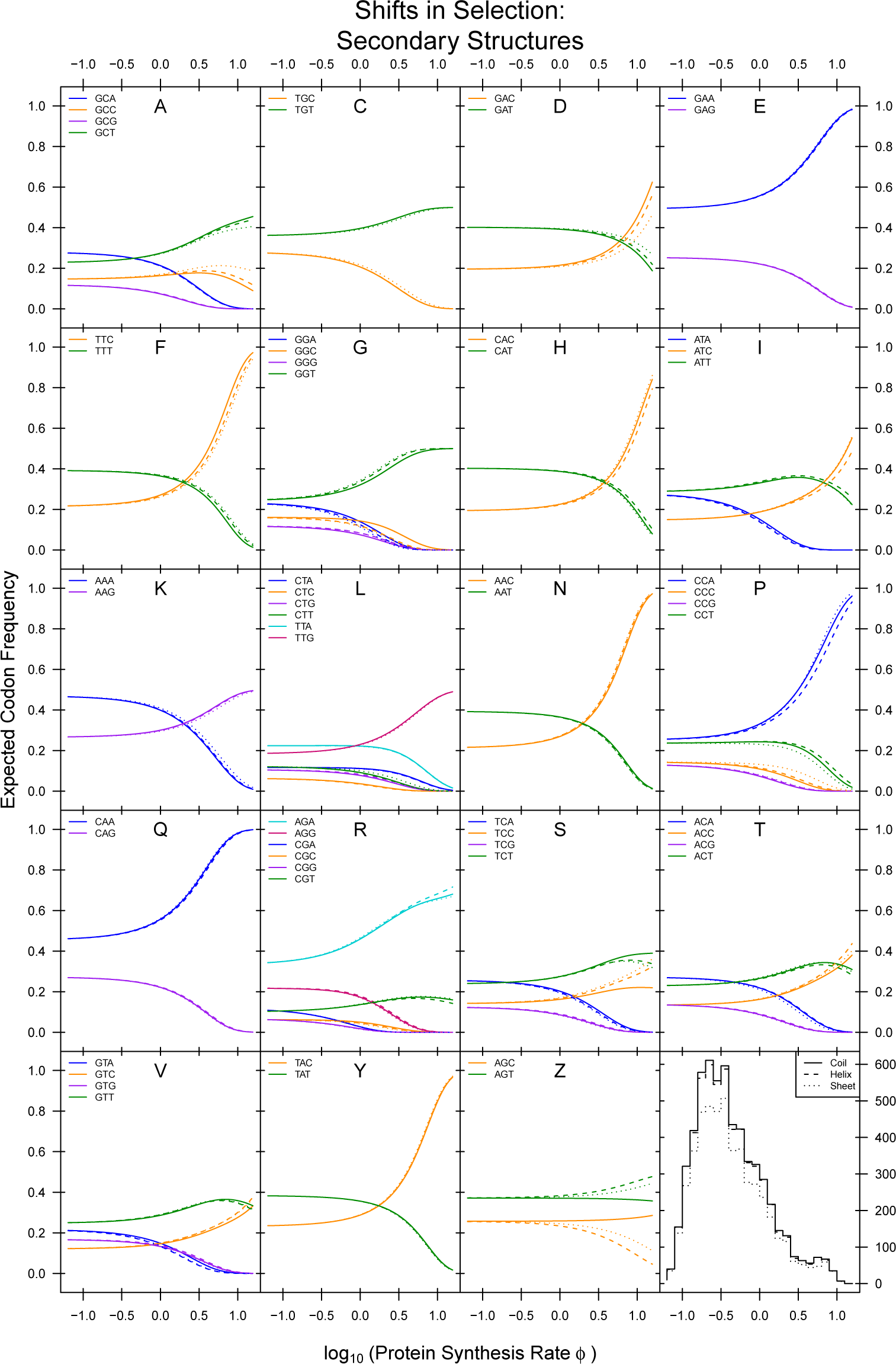
Comparison of expected codon frequencies across protein secondary structures as a function of protein production rates *φ*. Expected codon frequencies are estimated using equation 1 (see Materials and Methods). The bottom right histogram gives distributions of *φ*.

### Shifts in Selection: Secondary Structures

#### Intrinsically-disordered Regions Show Distinct Patterns of Selection on Codon Usage

In *S. cerevisiae*, we found that whether a region was structured or intrinsically-disordered better explained intragenic codon usage patterns than protein secondary structures in *S. cerevisiae* (Table 2, Models Y_5_ vs. Y_1_). Consistent with this, selection was 9% weaker, on average, in IDRs compared to structured regions (Deming Regression *β*^^^ = 0.905, 95% CI: 0.823 – 0.988, Figure S3A). In contrast, splitting codons into structured regions and IDRs in *E. coli* did a worse job of explaining intragenic codon usage patterns than secondary structures (Table 2, Model E_1_ vs. E_5_).

This was unsurprising given the rarity of IDRs in prokaryotic proteomes [49]. Despite being a worse fit compared to the secondary structure model in *E. coli*, the structured regions vs. IDR model is still a significant improvement over the null model (Tables 1 and 2, Model E_5_ vs. E_0_, ΔDIC = 129). Even though the Deming regression slope comparing structured regions and IDRs in *E. coli* was of similar magnitude to the same slope estimated for *S. cerevisiae*, it was not significant (Figure S3B, Deming Regression *β*^^^ = 0.933, 95% CI: 0.849 – 1.020).

Although selection on codon usage was weaker, on average, in IDRs of *S. cerevisiae*, we note a subset of amino acids demonstrate the opposite pattern in which selection against certain codons appears to be stronger: alanine (A), histidine (H), lysine (K), proline (P), and threonine (T) (Figure S3A). In addition, serine (S_4_ and S_2_) shows shifts in the selectively-favored codon in IDRs relative to structured regions, with the former showing preference for TCT and AGC. This is similar to the results observed in coils for *S. cerevisiae*, but in this case, the selective shifts clearly indicate one of the codons is preferred over the other (i.e. the 95% posterior probability intervals do not overlap with 0, Figure S3A). Alanine (A), serine (S_4_), and threonine (T) showed a similar pattern in *E. coli* (Figure S3B). Interestingly, many of these amino acids have a higher propensity for forming disordered regions and/or serve as sites for phosphorylation [50, 51]. Despite the apparent qualitative shift between structured regions IDRs in *S. cerevisiae*, as well as various quantitative shifts, these shifts have very little impact on the expected codon frequencies (Additional File 1, Figure S4).

We found that categorizing the predicted structured regions (from IUPRED2) based on the corresponding secondary structure predictions (from PsiPred) improved the overall model fits in both species (Models Y_6_ and E_6_), indicating differences in codon usage between secondary structures are not solely due to the presence of IDRs. Comparing selection estimates Δ*η* from the secondary structures (with IDRs removed) to Δ*η* estimated from IDRs suggests selection on codon usage is, on average, stronger in coil and sheet secondary structures compared to IDRs in *S. cerevisiae* (Additional File 1, Figure 3). Although a comparison of helices to IDRs has a slope estimate consistent with stronger selection in helices, this slope is not statistically significant (Deming regression *β*^^^ = 0.905, 95% CI: 0.806 – 1.000). In *E. coli*, all three Deming regression slopes are less than 1, but only the comparison between coils and IDRs is statistically significant (Figure S3F–H). When comparing Δ*η* between secondary structures after removing IDRs, we found that many codons still exhibited significant selective shifts between secondary structures (Figure 3). Notably, the observed selective shifts in *S. cerevisiae* on codons TCT (S_4_) and AGC (S_2_) in coils appears weakened or missing when IDRs are removed, suggesting the previously observed results were driven by differences in selection in IDRs.

**Figure 3:**
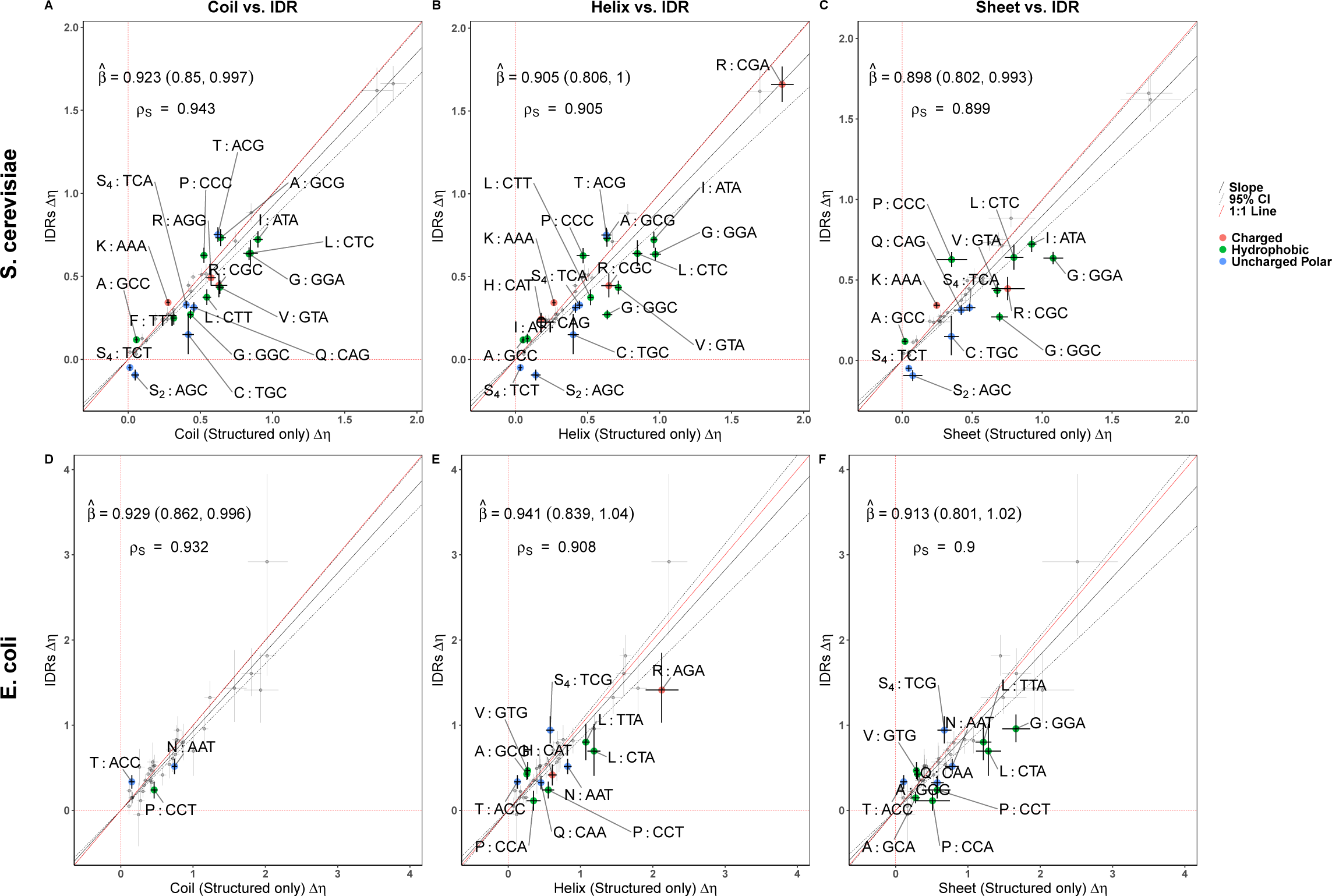
Comparison of selection estimates Δ*η* between secondary structures (predicted IDRs removed) and IDRs. Points represent Δ*η* values for each codon. Error bars represent the 95% posterior probability intervals. Codons showing significant selective shifts are col- ored by amino acid property. Negative values indicate a change in the selectively-favored codon relative to the genome-wide optimal. **(A,D)** Coils vs. IDRs. **(B,E)** Helices vs. IDRs. **(C,F)** Sheets vs. IDRs.

Given that all secondary structures were predicted to fall into structured regions or IDRs (Additional File 1, Table S3), we tested whether further dividing up the secondary structures into their corresponding structured and disordered components was better able to explain codon usage variation across regions. Seemingly the most logical split of coils into structured coils (i.e. those likely falling into protein domains) and IDR coils were better model fits than the models that relied solely upon secondary structure or disorder information in both *S. cerevisiae* and *E .coli* (Model Y_8_ and E_8_, respectively). In *S. cerevisiae*, splitting coils into the structured and disordered regions improved upon the model where the IDRs were taken from all three secondary structure classifications in (Model Y_8_ vs. Y_6_, ΔDIC = 34), but this model was a worse fit in *E. coli* (Model E_8_ vs. E_6_, ΔDIC = 28). Surprisingly, dividing both coils and helices into structured regions and IDRs in *S. cerevisiae* further improved the model fit (Model Y_10_ vs. Y_8_, ΔDIC = 83).

**Table 3:** Comparing models with termini (first and last 2 codons, respectively) of secondary structures separated from the core of the structure in *E. coli*. Results are for secondary structures of minimum length 6. H: helix. E: sheet. C: coil. ΔDIC = DIC*_i_ −* DIC_Best_.

| Structure Model | Secondary Structure | Groupings | | | $\Delta\text{DIC}$ |
| --- | --- | --- | --- | --- | --- |
|  |  | I | II | III |  |
| E<sub>1b</sub> | H | Termini | Core | – | 0 |
| E<sub>1c</sub> |  | N-terminus | Core | C-terminus | 13 |
| E<sub>1a</sub> |  | Whole Structure | – | – | 64 |
| E<sub>1e</sub> | E | Whole Structure | – | – | 0 |
| E<sub>1g</sub> |  | N-terminus | Core | C-terminus | 37 |
| E<sub>1f</sub> |  | Termini | Core | – | 43 |
| E<sub>1h</sub> | C | Whole Structure | – | – | 0 |
| E<sub>1i</sub> |  | Termini | Core | – | 6 |
| E<sub>1j</sub> |  | N-terminus | Core | C-terminus | 14 |

### Selection on codon usage varies at the termini of helices

In *S. cerevisiae*, we found no evidence that natural selection varies at the termini of sheets or coils based on model comparisons. This was regardless of our choice of minimum length of the structure or size of the termini (Additional File 1, Table S4 – S13). However, model comparisons did not allow us to rule out the possibility of differences in natural selection at the termini of helices. When restricting our analysis to helices with a minimum of 4 codons and defining the N and C-terminus to be the first and last two codons, respectively, we found the model allowing for differences at the termini improved upon the model assuming no differences at the termini by 14 DIC units (Additional File 1, Table S4). When helices were limited to those of length 6, the favored model did not allow for differences in selection between the core and the termini (Additional File 1, Table S6 – S7). We tested if there was a general difference in selection between helices of lengths 4 or 5 vs those with a length 6 or greater, but we found no support for this model over grouping all helices together (ΔDIC = 27). Considering helices of length 4 vs those with a length 5 or greater yielded similar results (ΔDIC = 30). We note that the effect weakened if excluding 3_10_-helices (DSSP G) and *π*-helices (DSSP I), but the overall patterns still suggests possible differences in selection at the termini of helices (Additional File 1, Table S14); however, we found no evidence to support general differences in selection on codon usage between these two helical structures and the traditional *α*-helices (ΔDIC = 17). Regardless, when comparing the Δ*η* estimates from the termini and the core of the helices, we find that 4 codons are significantly different: proline (P) codon CCC, glycine (G) codons GGA and GGC, and valine (V) codon GTT with the former (Additional File 1, Figure S7).

Model comparisons clearly indicate differences at termini relative to the core in *E. coli* in helical structures (Table 3 and Additional File 1, Tables S15 – S17). We note that this pattern is consistent if restricting the minimum length of helices to 4 – 7 codons. In this case, we find that 8 codons are significantly different between the termini and core (Additional File 1, Figure S6). On the other hand, sheets demonstrated variable patterns depending on the length. When restricting the length to a minimum of 4 or 5 codons, the model fits indicate there is a difference between the core and termini of sheets, but this disappears when restricting the analysis to only sheets of minimum length 6 or 7 codons. Although the same analysis in coils generally favored no differences in at the termini, we note that the ΔDIC scores were often less than 10 DIC units, meaning we could not confidently rule out differences in selection at the termini of coils.

### Interestingly, this effect disappeared when restricting the analysis to coils with at least 7 codons (Additional File 1, Table S17)

#### Previous claim of selective shifts at the start of helices is due to artifacts

Although we found support for some differences in selection at the termini of helices in *cerevisiae* (see Additional File 1, Figure S7 and Tables S6 – S16), we found no support for the model allowing for differences in selection at the second and third positions of helices [31] relative to the model assuming no differences in selection within helical structures (ΔDIC = 30) [31]. Using simulated data that assumes the strength and direction of selection for a codon is constant across the entire genome, we found the odds ratios reported by [31] were within the range of odds ratios generated using the simulated data (Figure 4). Similarly, patterns consistent with the enrichment of optimal codons at positions 1 and 4 are also observed in our simulated data. This suggests the enrichment of optimal and non-optimal codons at positions 1 and 4, and positions 2 and 3, respectively, of helices observed by [31] are an artifact of various confounding factors, such as amino acid biases and gene expression, that can shape codon usage patterns unrelated to natural selection.

**Figure 4:**
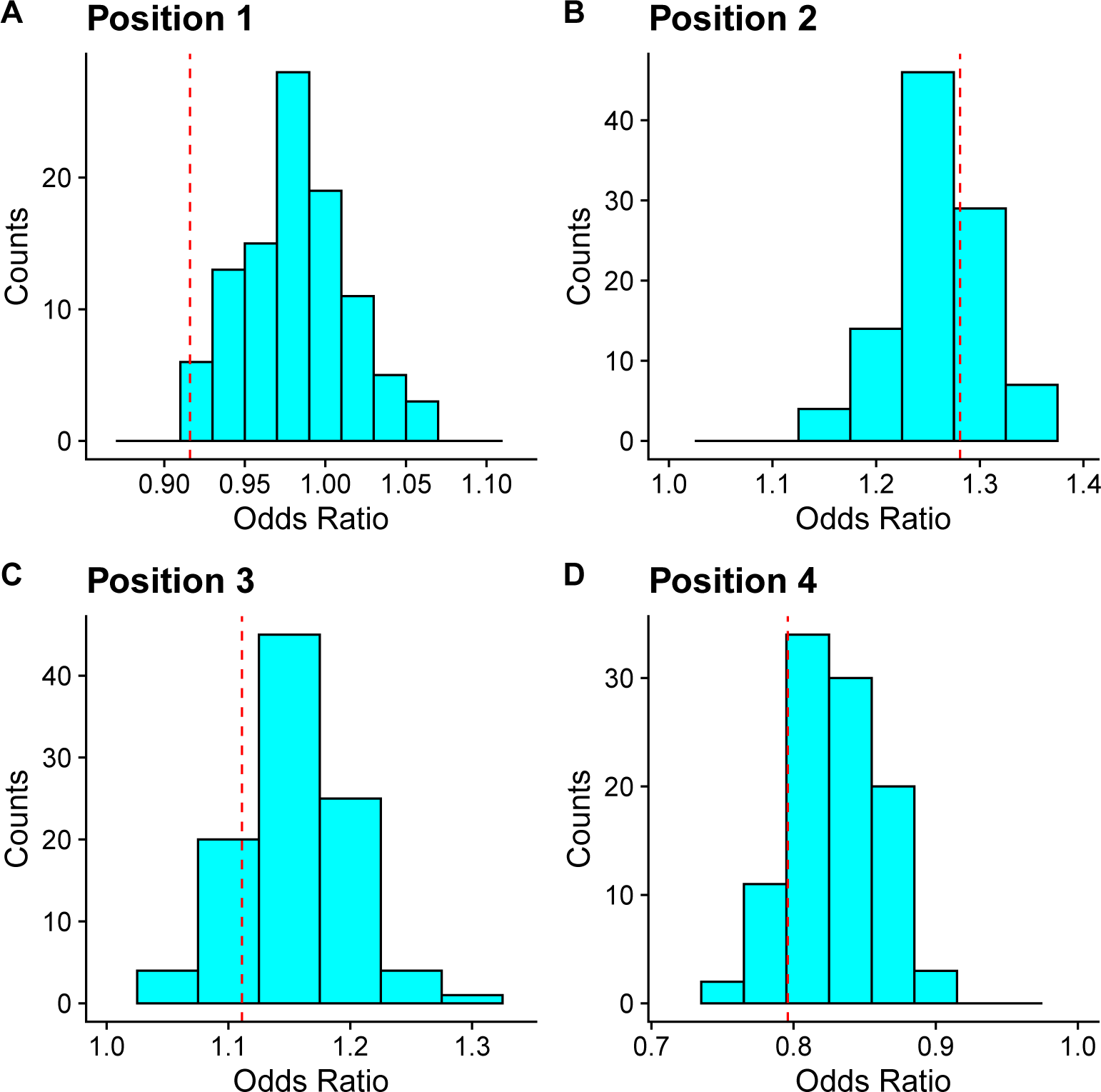
Odds ratio distributions for 100 simulated genomes by relative position in helices. Genomes were simulated assuming selection was identical across all positions in the helices. Dashed red lines indicate the odds ratio reported by [31] estimated from real codon usage patterns. Note that only proteins used in [31] were included in this analysis.

## Discussion

The goal of this work is to quantify the general relationship between codon usage and protein structure. This is in contrast to other work which has focused on identifying regions thought to be important to protein structure due to conservation of synonymous codon usage patterns between species [31, 37, 40]. To account for the effects of amino acid biases and gene expression, we used the population genetics-based model ROC-SEMPPR that explicitly includes the effects of mutation bias, selection, and genetic drift on synonymous codon usage patterns. Fitting ROC-SEMPPR to different genic regions allowed us to test for qualitative and quantitative differences in selection on codon usage related to protein structure in *S. cerevisiae* and *E. coli*. With one minor exception, i.e. the TCT and AGC codon pair in helices and sheets, we found no evidence for qualitative shifts in the nature of selection across protein secondary structures and IDRs in in either species. Instead, we observed small quantitative differences in the strength of selection between codon pairs across different protein structures. The contrasts with previous work which has claimed that certain protein structures show preferences for slow codons: overall, selection on codon usage is highly correlated between protein structures [36, 39]. So while our results are consistent with enrichment of a codon within a structure relative to another structure, our results do not indicate a qualitative change in codon preference between regions. We also find evidence that selection on codon usage may vary slightly at the termini of helical secondary structures in both *S. cerevisiae* and *E. coli*. Again, while our results are consistent with possible enrichment of non-optimal codons [32], our results do not indicate a qualitative change in codon preference at the termini of these structures.

The codon-specific nature of ROC-SEMPPR’s parameters Δ*η* allows the detection of codon-specific difference that may be hidden to other approaches that average over the codon and amino acid usage of a region. For example, our results were consistent with overall weaker selection on codon usage in IDRs, consistent with the findings of [38] and [39], but a subset of amino acids (alanine, glycine, histidine, lysine, proline, serine, and threonine) showed stronger selection between synonymous codons compared to structured regions in both species. Many of these amino acids have higher propensities for forming IDRs (e.g. proline, serine, lysine, alanine, glycine) or play common functional roles in IDRs such as serving as sites for phosphorylation (serine and threonine) [50, 51]. We speculate the apparent increased selection against certain codons in IDRs is due to stronger selection against missense errors for these amino acids in IDRs. Previous work found differences in codon usage between secondary structures were due to the inclusion of IDRs [39]. In contrast, many of the selective shifts between secondary structures we detected remained after removal of IDRs, although the magnitude of these differences did appear to be reduced. This suggests approaches relying on metrics such as CAI, which is calculated as an average of codon usage across all amino acids within a region, may obfuscate codon-specific selective shifts.

Perhaps the most surprising finding of this work was that splitting up both helices and coils based on disorder predictions provided the best overall model fit in *S. cerevisiae*. Although a “disordered coil” seems like a natural categorization for a protein structure, the phrase “disordered helix” or “disordered sheet” seems self-contradictory. Because previous work has found that IDRs can form transient secondary structures (particularly helices) under certain conditions [52, 53], this might explain some of the shift we see between “ordered” and “disordered” helicies and sheets. Clearly, further work in this area is needed.

Although we find some evidence of selective shifts at the termini of helical secondary structures in *S. cerevisiae*, we find no evidence supporting selective shifts at positions 2 and 3 of helical secondary structures [31]. The results in [31] is likely due to confounding factors that can also impact codon usage patterns, such as amino acid biases, which can be particularly problematic when using metrics such as tAI [16]. An important feature of our population genetics based approach is revealed from the null distributions generated under the assumption of no selective differences at positions 2 and 3. That is, the true null distribution of odds ratios is not centered around 1 as is usually assumed (Figure 4). These findings illustrate that while *ad-hoc* approaches to analyzing sequence data can be useful, care must be taken to ensure that the analysis is consistent with the corresponding evolutionary null model [19]. An evolutionary null model is often meant as the expected patterns if a trait were evolving exclusively in the absence of selection (i.e. genetic drift and mutational bias), but in our case, it refers to the patterns expected if the strength and direction of natural selection on codon usage were the same between two regions. Alternatively, population genetics approaches can be used for generating evolutionary null distributions for hypothesis testing with *ad-hoc* approaches, as we show here.

It is worth noting that the selectively-favored codons identified by ROC-SEMPPR may be different from those identified by other methods which fail to account for the effects of mutational biases. The optimal codons for each amino acid identified by ROC-SEMPPR and the tAI-based metric presented in [31] are in agreement for only 11 of the 19 amino acids (Additional File 1, Figure S8). For the remaining 8 amino acids, a common pattern emerges: the selectively-favored codon is also mutationally-favored based on ROC-SEMPPR’s parameter estimates, indicating that metric presented in [31] misidentifies the selectively-favored codon by not separating out the effects of mutational biases and genetic drift from selection.

Like all models, the biological realism of ROC-SEMPPR is sacrificed for the sake of tractability. Given the richness of biological systems, it would be interesting to build upon our analysis and take other factors that might affect codon usage bias into account. For example, recently [54] proposed alterations to elongation rates via mRNA secondary structure could modulate cotranslational protein folding, but previous work has also found that mRNA secondary structure rarely impacts ribosome elongation rates DelCampo2015. Conceivably, one could add mRNA stability as an additional category when defining different coding regions; however, this is beyond the scope of our focus.

Another evolutionary force that can shape codon usage is GC-biased gene conversion [55]. For the present study, we note that GC-biased gene conversion has previously shown to be present in yeast, but its impact is relatively small [56] and the effects of hitchhiking on codon usage in yeast have been somewhat controversial [57, 58, 59, 60]. For other organsisms whose genomes are believe to be more strongly impacted by GC-biased gene conversion, one could take a categorial approach similar to the one we use here and categorize genes by their recombination rate, if known. In theory, ROC-SEMPPR could be expanded to explicitly include GC-biased gene conversion as a quantiative term.

## Conclusions

We find that methods rooted in population genetics can be used to test for shifts in natural selection on codon usage. A key advantage of ROC-SEMPPR is it can be applied to any organism with a sequenced genome, requiring no other input, such as empirical estimates of gene expression [47, 44]. ROC-SEMPPR provides estimates of selection for individual codons, unlike other approaches based on heuristic measures of codon adaptation, such as CAI. We emphasize that we are attempting to quantify the average, genome-wide relationship between selection on codon usage and protein structure. These selective shifts are expected to reflect general mechanisms related to the folding of a protein structure [31, 39]. This is in a similar vein to work that has made broad statements about the preferences of a protein structure for certain codons, such as *α*-helices are preferentially encoded by translationally efficient codons [36]. Our work suggests that such statements are overly-simplistic, as the observed direction and magnitude of selective shifts clearly varies by codon, although these shifts are generally very small. A remaining challenge is to establish the relative importance of the the different selective forces that can shape the adaptive evolution of codon usage (e.g. translation efficiency, translation accuracy, mRNA secondary structure) related to protein structure. The direction of natural selection related to these aspects of codon usage do not always operate in the same direction [61]. Future work investigating differences in natural selection on codon usage related to protein folding, protein secretion, and other processes will benefit from the use of such models that directly capable of directly separating out the different selective forces shaping codon usage.

## Methods

Protein-coding sequences (CDS) and amino acids sequences for *S. cerevisiae* S288c (GCF 000146045.2) and *E. coli* K12 MG1655 (GCF 000005845.2) were downloaded from NCBI Refseq. Previous analysis of CUB in *E. coli* indicated approximately 750 genes had outlier codon usage patterns, many of which were hypothesized to be due to horizontal gene transfer [62].

Fitting these outlier genes with ROC-SEMPPR revealed selection on codon usage within these genes was anti-correlated with the remaining genes [42]. Here, our analysis of *E. coli* excludes these outlier genes.

### Identifying protein secondary structure

Our analysis makes use of both protein secondary structures determined empirically via methods like X-ray crystallography, and computationally via methods like PsiPred [48]. The empirical data is a more conservative dataset, with fewer proteins available but more accurate and reliable designations of protein secondary structures. The current implementation of PsiPred has an overall accuracy score of 84% [63], but secondary structure prediction algorithms generally struggle with accurately identifying the termini of secondary structures [32, 31]. Therefore, analyses of codon usage at secondary structure termini were based exclusively on empirically-determined secondary structures.

Empirically-determined protein secondary structures and corresponding protein sequences were obtained from the Protein Data Bank (PDB). Residues were grouped into three overarching structural groups based on their DSSP classification: helix (DSSP H, G, and I), sheet (DSSP E and B), and coil (DSSP S, T, and ’.’). This classification system is consistent with secondary structure prediction algorithms [48] and other analyses of codon usage patterns based on empirically-determined secondary structures [64, 36, 31, 32, 35]. Note that the classification symbol (.) is a catchall containing any amino acids not matching any other DSSP classifications. Protein sequences from PDB were aligned to the *S. cerevisiae* and *E. coli* proteomes using BLAST. Sequences were considered mapped to the proteomes if the PDB sequence covered 80% of the length of the protein and had a percent identity score of 95% or higher. This provided us with 1,097 and 1,285 protein sequences with empirically-determined secondary structures in *S. cerevisiae* and *E. coli*, respectively. This dataset was used for comparing selection on codon usage between and within secondary structures.

Protein secondary structures were predicted for all nuclear protein sequences for *S. cerevisiae* and for 1,742 proteins in *E. coli* using the PsiPred software [48] at default settings. PsiPred combines the secondary structural classifications of DSSP into helices (H), sheets (E), and coils (C).

### Identifying structured and intrinsically-disordered regions

Unlike protein secondary structures, empirically-determined intrinsically-disordered regions (IDRs) are rare. The DisProt database includes only 134 proteins with IDRs for *S. cerevisiae*. Thus, our analysis of codon usage patterns in IDRs and structured regions in *S. cerevisiae* and *E. coli* relied on predicted IDRs using IUPRED2 [65], which provides a quasi-probability of the an amino acid falling into a disordered region, using default settings. An amino acid with a quasi-probability of greater than 0.5 is more likely to be disordered, while a quasi-probability less than 0.5 is more likely to be structured; thus, amino acids with a score less than or equal to 0.5 were classified as structured, while amino acids with a score greater than 0.5 were classified as disordered, consistent with the analysis done by [39].

### Analysis with ROC-SEMPPR

All analyses of CUB was performed using ROC-SEMPPR with the R package AnaCoDa [47]. We note ROC-SEMPPR assumes weak selection. To meet this assumption, serine was split into separate codon groups: the 4 codon group TCN (S_4_) and the 2 codon group AGN (S_2_).

For any amino acid with *n_aa_* synonymous codons, the probability of observing codon *i* in gene *g* can be described by the equation

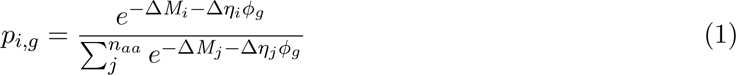

where Δ*M* represents mutation bias, Δ*η* represents natural selection, and *φ* represents the evolutionary average protein production rate of gene *g*. Note that Δ indicates the mutation bias and natural selection parameters are relative to a reference codon. ROC-SEMPPR’s mutation bias Δ*M* parameter represents the log of the ratio of mutation rates between two synonymous codons [44]. Although originally described as being proportional to relative differences in translation efficiency between two synonymous codons [44], Δ*η* can also be interpreted as the critical population genetics parameter *sN_e_*, where *N_e_* is the effective population size and *s* represents the selection coefficient relative to the reference codon, here chosen to be the most selectively favored codon for an amino acid. Because ROC-SEMPPR assumes the strength of selection varies with a gene’s expression level *φ* and scales this term such that the average level of expression across genes is *φ* = 1, Δ*η* represents the strength and direction of natural selection for a codon in a gene with an average expression level. For genes with lower or higher expression than average, the strength of this selection simply scales with *φ*, i.e. *sN_e_* = Δ*ηφ*.

A deeper understanding of the model parameters can be obtained by considering the cases of no protein production *φ* = 0 and average protein production *φ* = 1. We note that ROC-SEMPPR scales *φ* such that the *E*[*φ*] = 1. In the case of no protein production, natural selection on codon usage is completely absent, resulting in mutation biases determining the synonymous codon frequencies. In case of average protein production, the synonymous codon frequencies will reflect the relative strengths and directions of mutation bias and natural selection (proportional to drift). For example, if the mutation bias is stronger and in the opposite direction of natural selection (i.e. mutation and selection favor different codons), then the mutationally-favored codon is expected to be more frequent in an average expression gene. Importantly, *φ* scales the strength of natural selection such that strong mutation biases can be (but not necessarily will be) overwhelmed in highly expressed genes. Previous work indicates ROC-SEMPPR’s parameter estimates correlate well with empirical measurements in *S. cerevisiae* and *E. coli* [44, 42].

### Analysis of selective shifts on codon usage between protein structures

ROC-SEMPPR was fit to all protein-coding sequences in *S. cerevisiae* and *E. coli* to obtain gene-specific estimates of protein production rates *φ* and codon-specific estimates of mutation bias Δ*M* . Protein-coding sequences were then partitioned based on the codons corresponding secondary structure (based on empirically-determined or predicted structures). To determine if codon usage patterns are statistically different between protein secondary structures, structural groupings were combined, e.g. helices and sheets were combined into one group as opposed to treating them as separate groups. These structural groupings were then further merged into different models such that each structure category was represented once, either as a standalone group (e.g. helix) or grouped with another secondary structure (e.g. helix and sheet as one group). To be clear, “different models” simply refers to different ways in which different secondary structures were grouped. This ensured the sequence data (i.e. the number of codons, protein-coding sequences, etc.) is the same across all models, making them directly comparable. We note that the first 35 codons of all genes were excluded to help reduce the impact of a weaker selection on codon usage at the 5’-end of genes [66].

A similar analysis to the one outlined above for comparing codon usage between secondary structures was performed based on the predictions using IUPRED2, in which we compared a model which had structured and disordered regions as separate groupings to a model which treated them as one grouping. Finally, an analysis was performed which combined information from PsiPred and IUPRED2 to classify amino acids based on both methods for classifying structural information. This allowed us to distinguish coils which may be found as part of a larger structured domain from coils part of intrinsically-disordered regions.

### Analysis of selective shifts on codon usage within protein structures

To examine variation in codon usage within secondary structures, empirically-determined secondary structures were divided into the N-terminus and C-terminus regions, with all codons in between being classified as the core of the secondary structures. To assess robustness of our results, we varied the minimum number of amino acids for a secondary structure between 4 and 10 and varied the termini length to be 2 or 3 amino acids. To test the hypothesis presented by [31] in *S. cerevisiae*, helices of minimum length 6 amino acids were split up into the second and third codons (relative to the start of the helix) and the remainder of the helix.

### Comparing model fits and estimates of selection

For statistically comparing codon usage patterns, ROC-SEMPPR model fits were compared using the Deviance Information Criterion (DIC) [67]. Briefly, DIC is a Bayesian information criterion which tries to balance the overall model fit to the data as determined by the posterior distribution and the number of parameters used to fit the data. If the level or nature of selection on codon usage differs between two structures, then it is expected a model treating these structures as separate groupings will have a better (lower) DIC score than model fits treating the structures as single (or merged) groupings. We follow the general rules of thumb for comparing models using information criterion [68]. A model that differs from the minimum DIC model by fewer than 2 DIC units has substantial statistical support. A difference in the range of the 2-4 DIC units are still considered to have strong support, while a difference of 4-7 DIC units are considered to have less support. However, a model that differs from the minimum DIC model by 10 or more DIC units can generally be disregarded. We note that all ΔDIC values will represent DIC*_i_ −* DIC_Best_, where DIC*_i_* is the DIC score of the *i*^th^ model and DIC_Best_ is the minimum DIC score (i.e. the best model) of the models under consideration. This means that all reported ΔDIC values will fall into the range [0*, ∞*).

Comparing models via DIC indicates differences in selection on codon usage between structural regions, but does not tell us how they differ. Similar to [42], we broadly compared codon-specific estimates of selection Δ*η* between structural groupings using a model-II regression, which accounts for errors in both the independent and dependent variables [69]. In this work, we used the Deming Regression, as implemented in the R package **deming**. A Deming regression on Δ*η* estimated from different structural grouping with a slope of *β*^^^ = 1 (i.e. *y* = *x*) would suggest there is not a general shift in natural selection on codon usage between the two groupings that can be described by the functional relationship Δ*η_B_* = *β*Δ*η_A_*. On the other hand, a Deming regression slope *β*^^^ significantly different from 1 is consistent with an overall shift in natural selection on codon usage between the two structural groupings being compared. For each amino acid, its corresponding Δ*η* values were scaled relative to its optimal synonymous codon, i.e. the one most favored by natural selection based on fitting the null model where selection on codon usage does not vary across structural categories; specifically, models *Y*_0_ and *E*_0_ for *S. cerevisiae* and *E. coli*, respectively. As a result, the null model optimal codon Δ*η* value is always 0 and its synonyms are always Δ*η >* 0, unless there’s a structure-specific shift in the optimal codon. In this case, there can be Δ*η* values less than 0. Importantly, the Deming regression only summarizes a possible overall shift in the strength or direction of selection on codon usage between two structural groupings, but does not rule out the possibility that selection is different between specific codons. This information can be obtained by comparing the Δ*η* estimates for a codon across the different protein structures. We focus on the codons for which the Δ*η* 95% posterior probability intervals do not overlap between structural groupings, as we are most confident in the sign of any shift in selection.

### Simulating codon usage patterns for model validation

To test if we are able to detect shifts in selection across protein regions, we simulated codon usage of two regions under the ROC-SEMPPR model using the empirically determined helices and coils in *S. cerevisiae* (1,097 genes represented) as templates. In one region, denoted as Region A, all codons are evolving under the same selective pressure. In the other region, denoted as Region B, some percentage of codons (1%, 10%, 50%, 100%) were randomly selected to be under the opposite selective pressure of Region A, with the remaining codons under the same selective pressure as Region A. This approach allows us to assess how the percentage of codons within a region To make sure our approach is robust to factors such as protein structure-specific amino acid biases, we simulated approximately 6,000 *S. cerevisiae* genomes such that the selective pressure was the same across all protein structures (i.e. all structure had the same values of selection parameter Δ*η*). Codons from the simulated dataset were assigned to different structures based the computationally predicted structures from PsiPred or IUPRED2. Qualitative and quantitative shifts between helices, sheets, and coils, and between structured regions and IDRs were determined as described in *Comparing model fits and estimates of selection*.

### Evaluating effects of confounding factors in analyses of position-specific codon usage

Previous work found that positions 2 and 3 (relative to the start) of helices in *S. cerevisiae* were enriched in non-optimal codons [31]. To determine if this pattern can be generated by amino acid biases or gene expression, the yeast genome was independently simulated 100 times using **AnaCoDa** [47] under the ROC-SEMPPR model with the genome-wide estimates of selection Δ*η* and mutation bias Δ*M* , as well as the gene-specific estimates of protein production rates *φ*, such that the nature of selection was the same for every codon across the genome. Proteins used by [31] for their analysis of position-specific codon usage were pulled and all helices were aligned by position. For each position, enrichment of non-optimal codons based on the nTE metric (as defined in [31]) was tested using a Fisher’s exact test. This generated a distribution of 100 odds ratios per position, which were then compared to the reported odds ratios in [31].

## Abbreviations

CUB: Codon Usage Bias
ROC-SEMPPR: Ribosome Overcost Stochastic Evolutionary Model of Protein Production Rates
IDR: Intrinsically Disordered Regions
DIC: Deviance Information Criterion
CAI: Codon Adaptation Index
tAI: tRNA Adaptation Index

## Availability of data and materials

Protein-coding sequences for *S. cerevisiae* and *E. coli* are available from National Center for Biotechnology Information (NCBI) RefSeq (https://ftp.ncbi.nlm.nih.gov/genomes/refseq/ fungi/Saccharomyces_cerevisiae/latest_assembly_versions/GCF_000146045.2_R64/ and https://ftp.ncbi.nlm.nih.gov/genomes/all/GCF/000/005/845/GCF_000005845.2_ASM584v2/, respectively). All other data and custom scripts are provided via the Github repository ( https://github.com/acope3/CUB_Protein_Structure_Analysis). The empirical secondary structures for *S. cerevisiae* and *E. coli* were pulled from a file downloaded from the Protein Data Bank (PDB), but due to a change in the API PDB, this file appears to be no longer available.

Due to the size of the original files downloaded from PDB, they are not included in the Github repository (although the relevant files for the organisms are included), but are available upon request.

## Declarations

### Ethics approval and consent to participate

Not applicable.

### Consent for publication

Not applicable.

### Authors’ contributions

A.L.C performed all analyses described in this work. M.A.G and A.L.C both contributed to the study design, interpretation of data, and writing of the manuscript. Both authors read and approved the final manuscript.

## Acknowledgements

The authors would like to thank John Favate and for his helpful comments in constructing this manuscript.

## Authors’ information

Current institution for A.L.C: Rutgers University.

## Additional Files

Additional File 1 — Supplemental Figures/Tables

PDF also includes supplemental figures and tables referenced in the text.

