## Supplemental Material for "Quantifying shifts in natural selection on codon usage between protein regions: A population genetics approach"


Wednesday 8th September, 2021

### Supplemental Figures

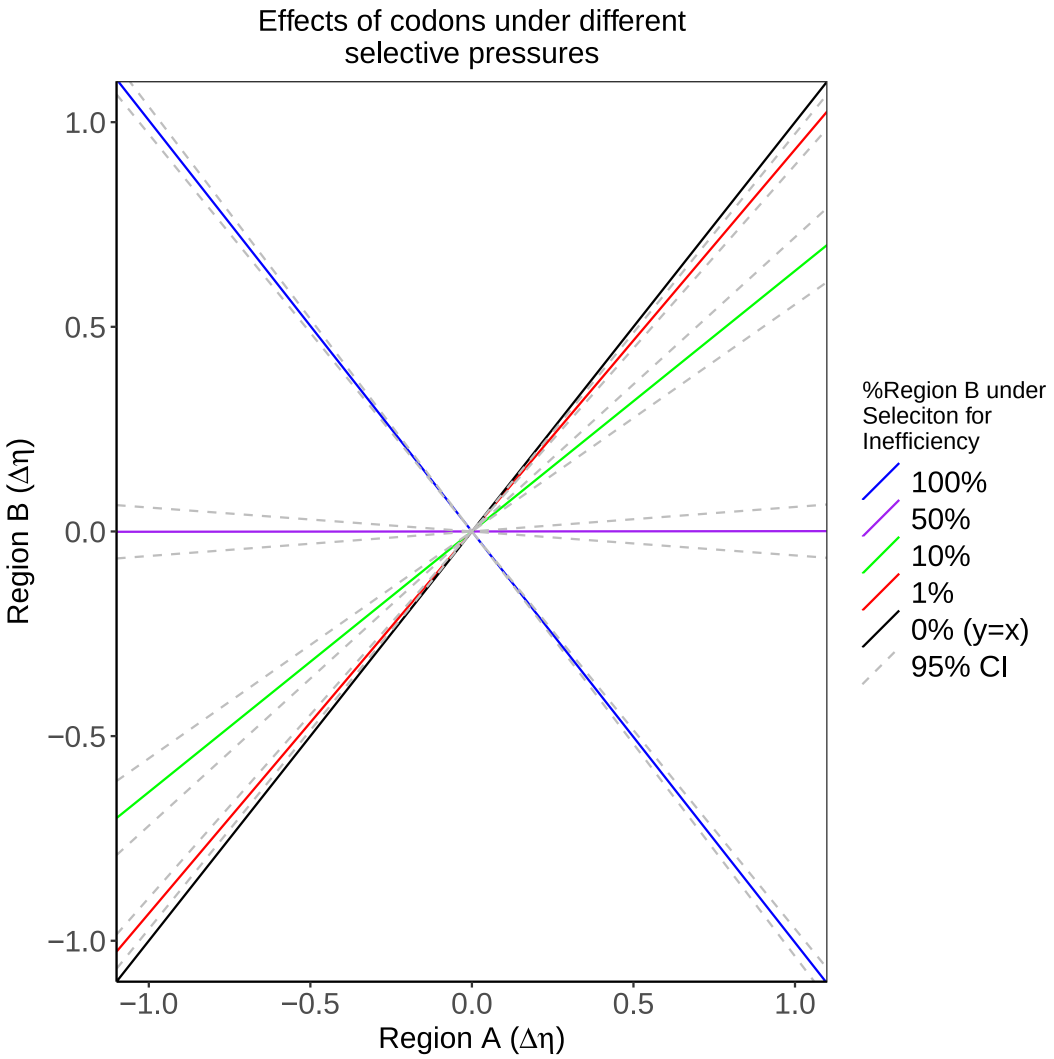


Figure S1: Comparing natural selection $\Delta\eta$ across simulated data with varying percentage of codons in Region B under selection for translation inefficiency. Note that this means these codons have $\Delta\eta$ values perfectly anti-correlated with the $\Delta\eta$ for the rest of the simulated genes. When 50% of Region B is under selection for translation inefficiency with the remaining codons under selection against inefficiency, ROC-SEMPPR is unable to determine the selectively favored codon.

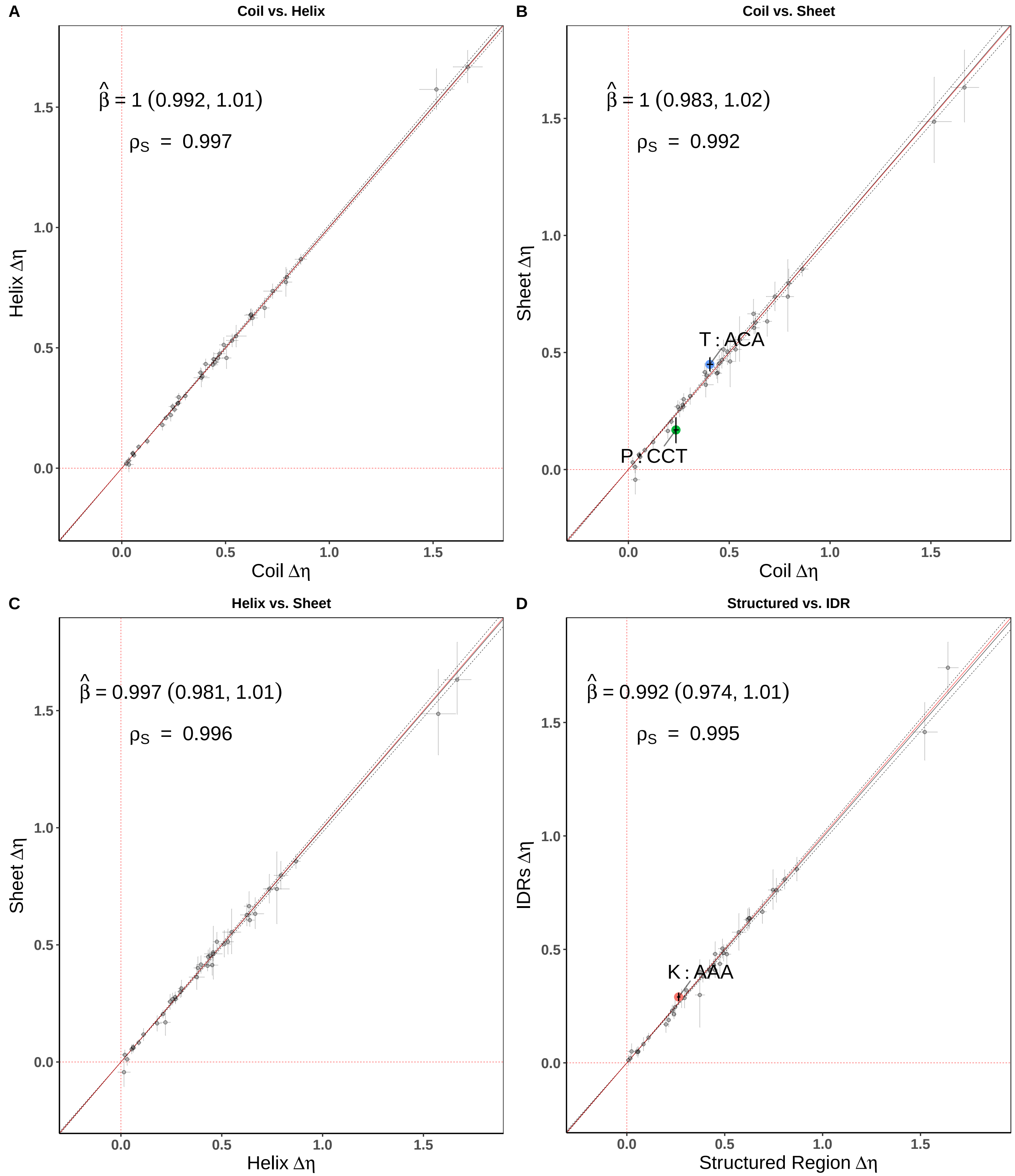


Figure S2: Comparison of mean-centered $\Delta\eta$ estimated from simulated codon usage data for different structures using *S. cerevisiae* protein-coding . **(A)** Coil vs. Helix. **(B)** Coil vs. Sheet. **(C)** Helix vs. Sheet. **(D)** Structured vs. IDRs

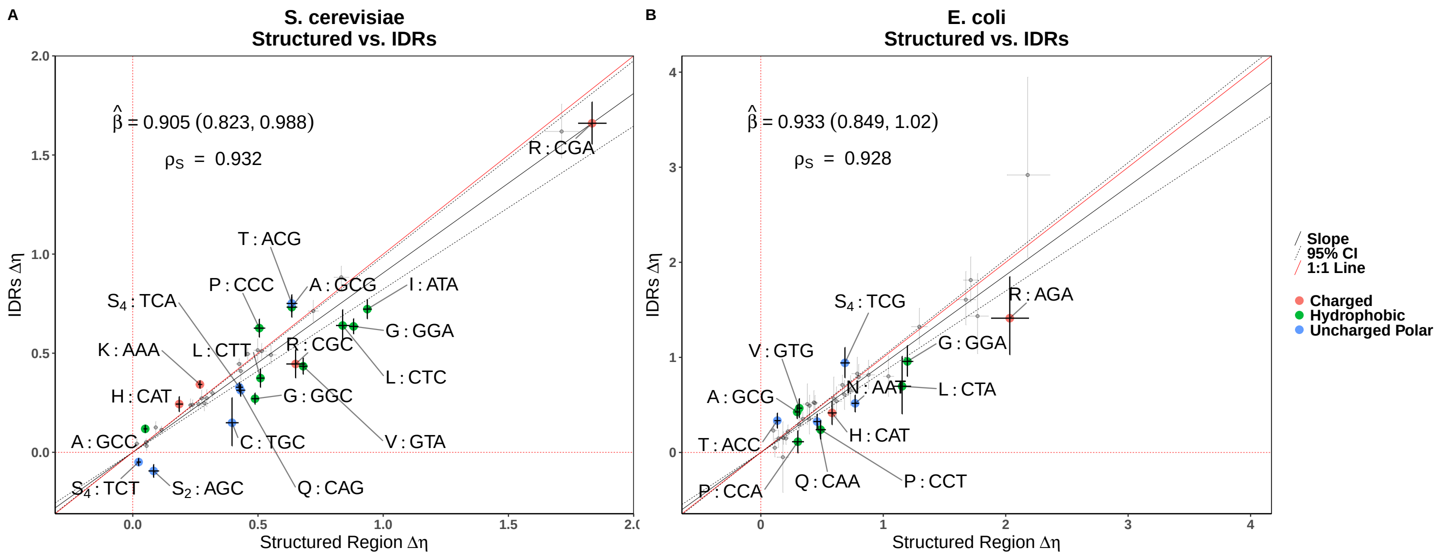


Figure S3: Comparison of selection estimates $\Delta\eta$ between structured regions and IDRs. Points represent $\Delta\eta$ values for each codon. Error bars represent the 95% posterior probability intervals. Codons showing significant selective shifts are colored by amino acid property. Negative values indicate a change in the selectively-favored codon relative to the genome-wide optimal. **(A,B)** Structured vs. IDRs.

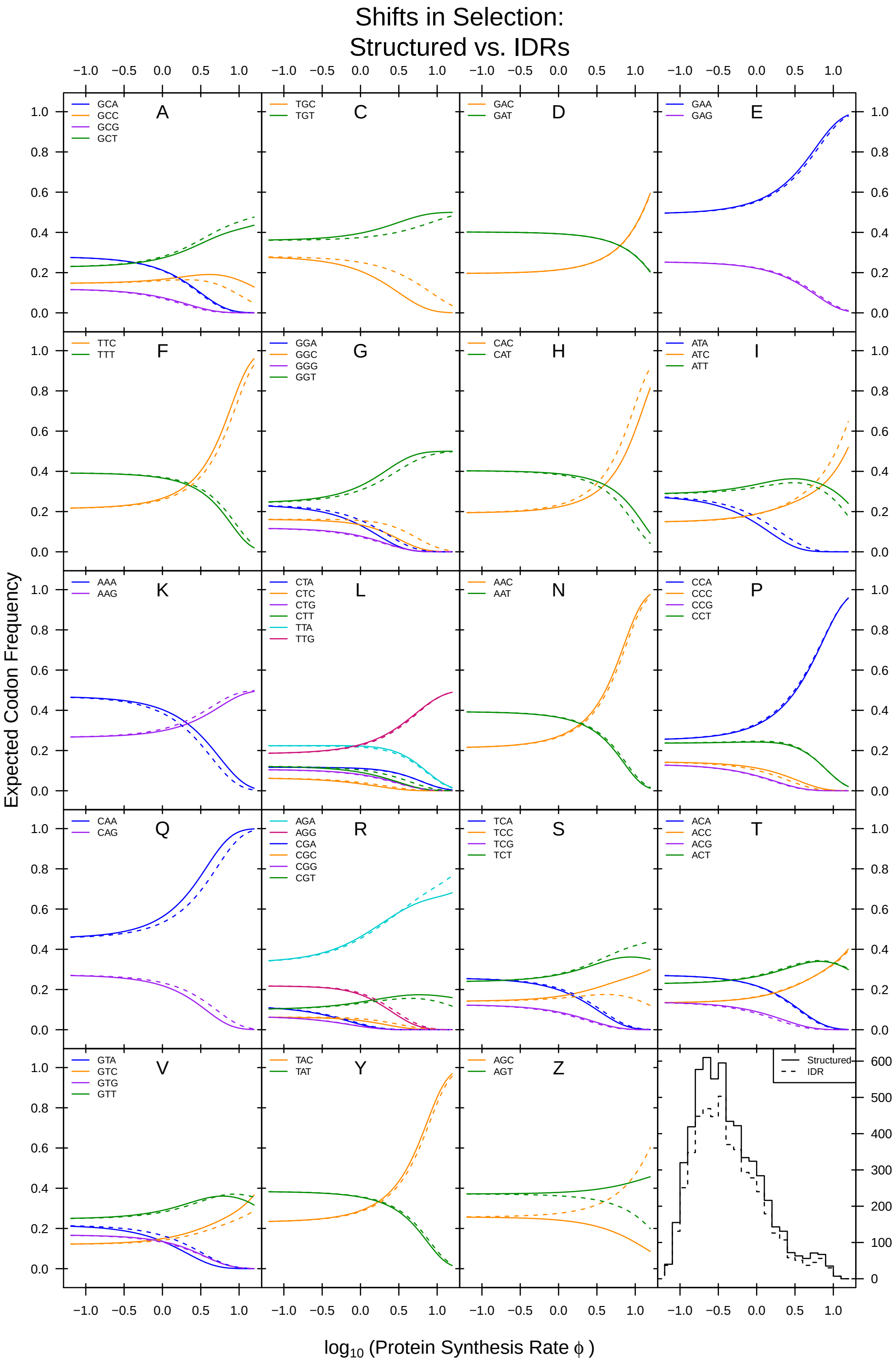


Figure S4: Comparison of expected codon frequencies in structured regions and IDRs as a function of protein production rates $\phi$. Expected codon frequencies are estimated using equation 1 (see Materials and Methods).

.

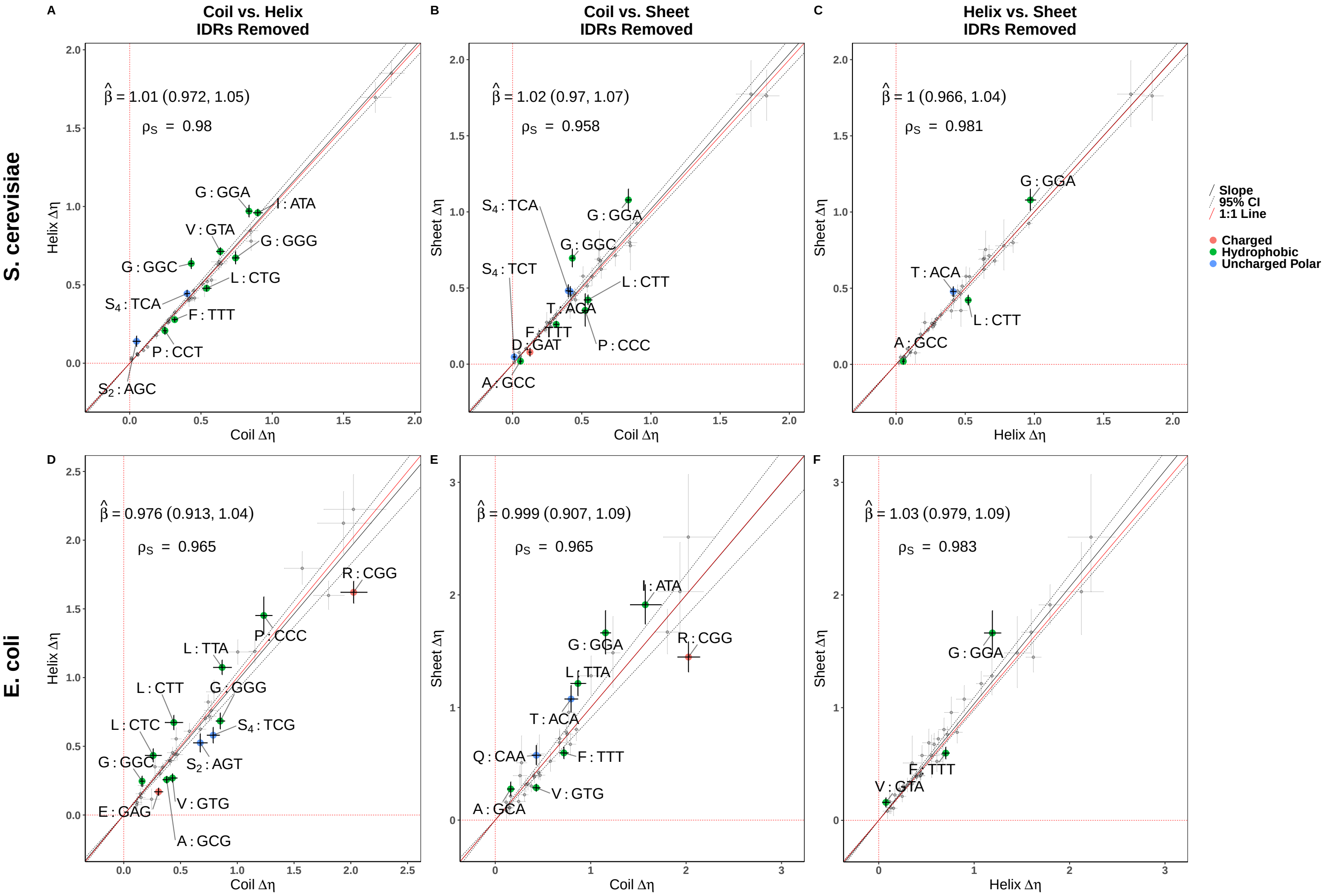


Figure S5: Comparison of selection estimates $\Delta\eta$ between different protein secondary structures with IDRs removed. $\Delta\eta$ is rescaled relative to the genome-wide optimal codon. Points represent $\Delta\eta$ values for each codon, while error bars represent the 95% posterior probability intervals. Codons showing significant shifts in selection are colored by amino acid property. Negative values indicate a change in the selectively-favored codon relative to the genome-wide optimal. The Deming regression slope and 95% confidence intervals are represented by solid and dashed black lines, respectively. **(A,D)** Coils vs. Helices. **(B,E)** Coils vs. Sheets. **(D,F)** Helices vs. Sheets.

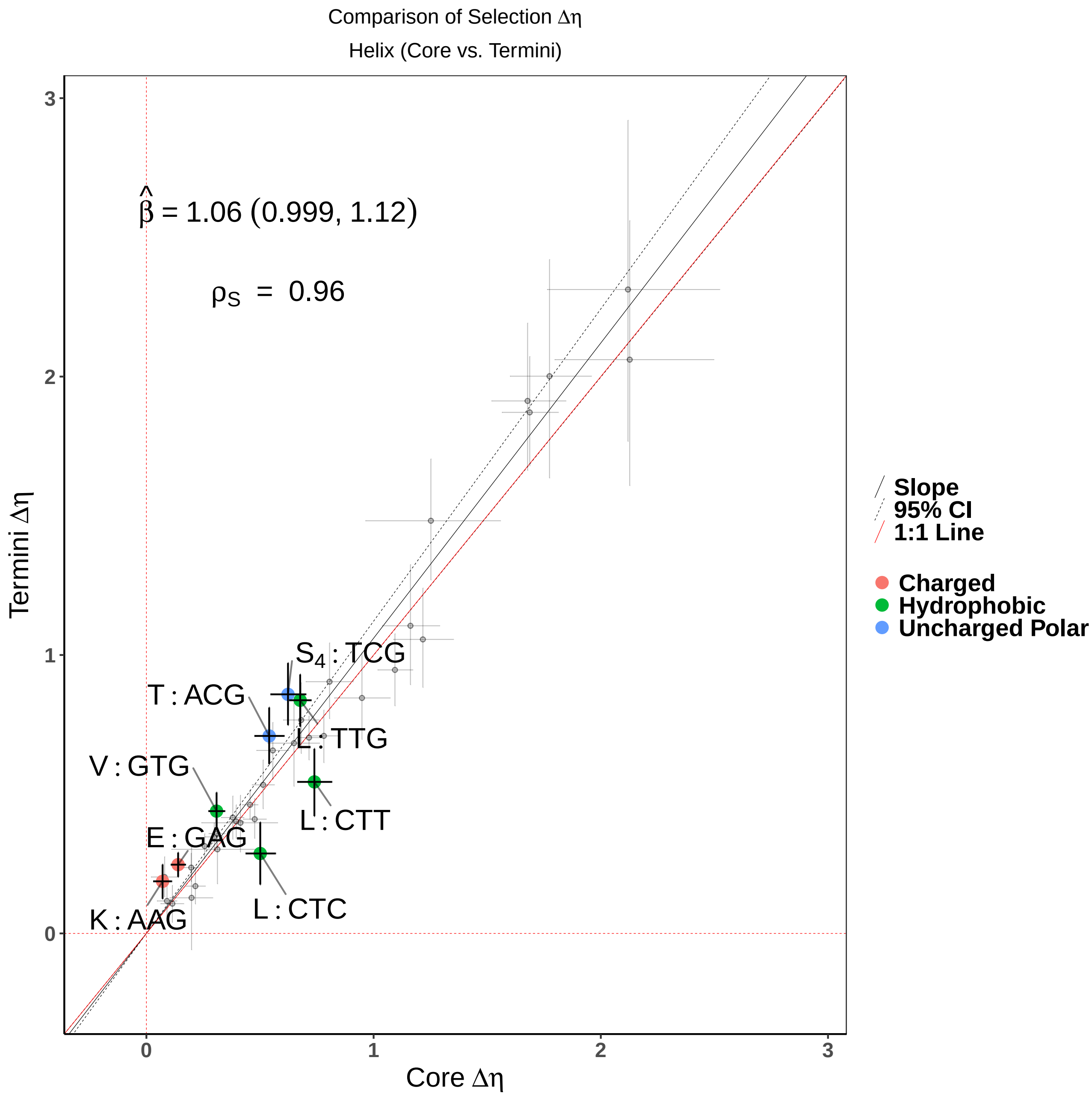


Figure S6: Comparison of selection $\Delta\eta$ between the termini and the core of helices in *E. coli*. Data was restricted to helices of minimum length 6 and treating the termini as the first and last 2 codons.

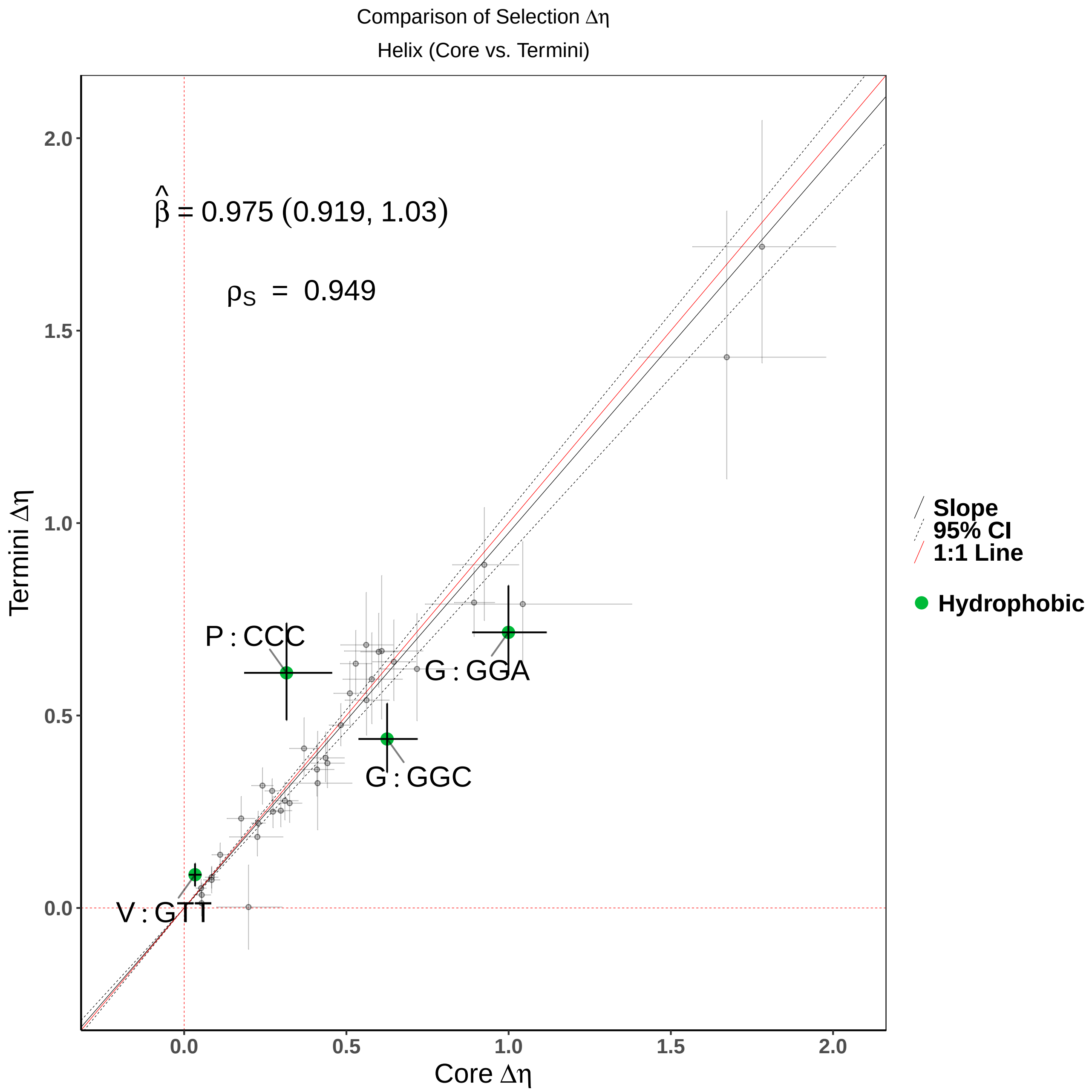


Figure S7: Comparison of selection $\Delta\eta$ between the termini and the core of helices. Data was restricted to helices of minimum length 4 and treating the termini as the first and last 2 codons.

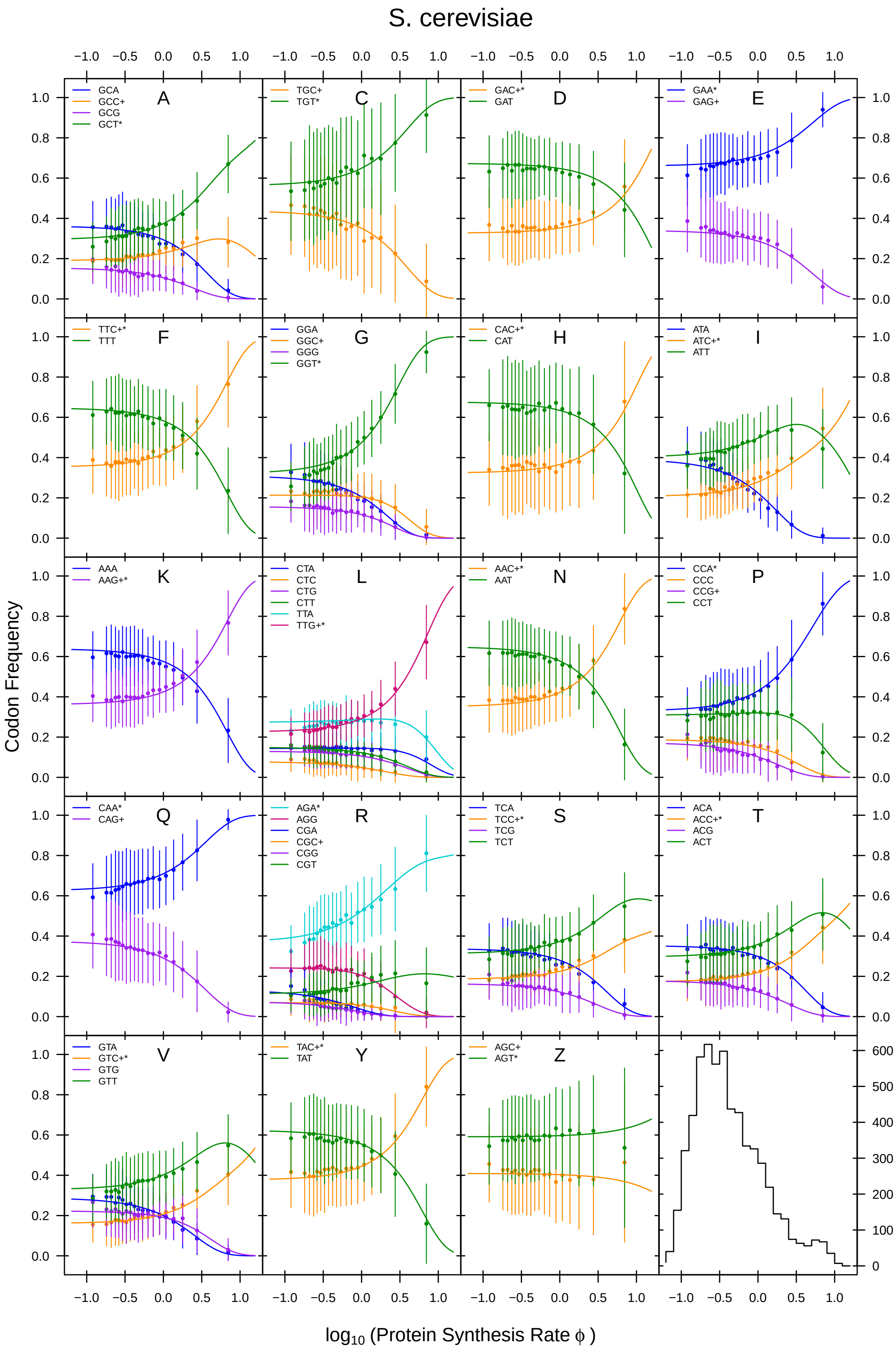


Figure S8: Observed (points) and expected (lines) codon frequencies for each amino acid as a function of protein synthesis (production) rates $\phi$, as well as the distribution of $\phi$ for all genes. Note that serine is split into two amino acid groups, S = $S_4$ and Z = $S_2$. An * indicates the preferred codon identified by ROC-SEMPPR, while + indicates the preferred codon identified by nTE.

### Supplemental Tables

Table S1: Comparison of model fits examining variation in codon usage in *S. cerevisiae* categorized using empirically-determined protein secondary structures from PDB. Models with a DIC score greater than the null model (no difference in codon usage between secondary structures) were excluded from the table. H: helix. E: sheet. C: coil.

| Structure Model | Groupings I | II | III | ΔDIC |
|---|---|---|---|---|
| $Y_1$ | H | E | C | 0.00 |
| $Y_2$ | HE | – | C | 1 |
| $Y_4$ | HC | E | – | 21 |
| $Y_0$ | HEC | – | – | 34 |

Table S2: Comparison of model fits examining variation in codon usage in *E. coli* categorized using empirically-determined protein secondary structures from PDB. Models with a DIC score greater than the null model (no difference in codon usage between secondary structures) were excluded from the table. H: helix. E: sheet. C: coil.

| Structure Model | Groupings I | II | III | ΔDIC |
|---|---|---|---|---|
| $E_1$ | H | E | C | 0.00 |
| $E_2$ | HE | – | C | 20 |
| $E_3$ | H | EC | – | 90 |
| $E_4$ | HC | E | – | 101 |
| $E_0$ | HEC | – | – | 145 |

Table S3: Breakdown of predictions by PsiPred (secondary structures) and IUPRED2 (disordered) in *S. cerevisiae*. Each value represents the number of amino acids falling into the corresponding categories. The percentages are relative to total number of amino acids predicted to be structured and disordered.

| | Structured | Disordered |
|---|---|---|
| Coil | 1,015,435 (0.43) | 451,974 (0.79) |
| $\alpha$-helix | 1,062,128 (0.45) | 106,645 (0.19) |
| $\beta$-sheet | 281,425 (0.12) | 13,899 (0.02) |

Table S4: Comparing models with termini of secondary structures separated from the core of the structure only considering secondary structures of 4 codons or longer with the terminus defined as the first and last 2 codons. H: helix. E: sheet. C: coil.

| Structure Model | Secondary Structure | Groupings I | II | III | ΔDIC |
|---|---|---|---|---|---|
| $Y_{1b}$ | H | Termini | Core | – | 0 |
| $Y_{1a}$ | | Whole Structure | – | – | 14 |
| $Y_{1c}$ | | N-terminus | Core | C-terminus | 20 |
| $Y_{1d}$ | E | Whole Structure | – | – | 0 |
| $Y_{1e}$ | | Termini | Core | – | 39 |
| $Y_{1f}$ | | N-terminus | Core | C-terminus | 75 |
| $Y_{1g}$ | C | Whole Structure | – | – | 0 |
| $Y_{1h}$ | | Termini | Core | – | 38 |
| $Y_{1i}$ | | N-terminus | Core | C-terminus | 82 |

Table S5: Comparing models with termini of secondary structures separated from the core of the structure only considering secondary structures of 5 codons or longer with the terminus defined as the first and last 2 codons. H: helix. E: sheet. C: coil.

| Structure Model | Secondary Structure | Groupings I | II | III | ΔDIC |
|---|---|---|---|---|---|
| $Y_{1b}$ | H | Termini | Core | – | 0 |
| $Y_{1a}$ | | Whole Structure | – | – | 3 |
| $Y_{1c}$ | | N-terminus | Core | C-terminus | 27 |
| $Y_{1d}$ | E | Whole Structure | – | – | 0 |
| $Y_{1e}$ | | Termini | Core | – | 37 |
| $Y_{1f}$ | | N-terminus | Core | C-terminus | 73 |
| $Y_{1g}$ | C | Whole Structure | – | – | 0 |
| $Y_{1h}$ | | Termini | Core | – | 39 |
| $Y_{1i}$ | | N-terminus | Core | C-terminus | 86 |

Table S6: Comparing models with termini of secondary structures separated from the core of the structure only considering secondary structures of 6 codons or longer with the terminus defined as the first and last 2 codons. H: helix. E: sheet. C: coil.

| Structure Model | Secondary Structure | Groupings I | II | III | ΔDIC |
|---|---|---|---|---|---|
| $Y_{1a}$ | | Whole Structure | – | – | 0 |
| $Y_{1b}$ | H | Termini | Core | – | 2 |
| $Y_{1c}$ | | N-terminus | Core | C-terminus | 33 |
| $Y_{1d}$ | | Whole Structure | – | – | 0 |
| $Y_{1e}$ | E | Termini | Core | – | 33 |
| $Y_{1f}$ | | N-terminus | Core | C-terminus | 57 |
| $Y_{1g}$ | | Whole Structure | – | – | 0 |
| $Y_{1h}$ | C | Termini | Core | – | 47 |
| $Y_{1i}$ | | N-terminus | Core | C-terminus | 96 |

Table S7: Comparing models with termini of secondary structures separated from the core of the structure only considering secondary structures of 6 codons or longer with the terminus defined as the first and last 3 codons. H: helix. E: sheet. C: coil.

| Structure Model | Secondary Structure | Groupings I | II | III | ΔDIC |
|---|---|---|---|---|---|
| $Y_{1a}$ | | Whole Structure | – | – | 0 |
| $Y_{1b}$ | H | Termini | Core | – | 3 |
| $Y_{1c}$ | | N-terminus | Core | C-terminus | 26 |
| $Y_{1d}$ | | Whole Structure | – | – | 0 |
| $Y_{1e}$ | E | Termini | Core | – | 46 |
| $Y_{1f}$ | | N-terminus | Core | C-terminus | 86 |
| $Y_{1g}$ | | Whole Structure | – | – | 0 |
| $Y_{1h}$ | C | Termini | Core | – | 42 |
| $Y_{1i}$ | | N-terminus | Core | C-terminus | 94 |

Table S8: Comparing models with termini of secondary structures separated from the core of the structure only considering secondary structures of 7 codons or longer with the terminus defined as the first and last 2 codons of the structure. H: helix. E: sheet. C: coil.

| Structure Model | Secondary Structure | Groupings I | II | III | ΔDIC |
|---|---|---|---|---|---|
| $Y_{1a}$ | | Whole Structure | – | – | 0 |
| $Y_{1b}$ | H | Termini | Core | – | 14 |
| $Y_{1c}$ | | N-terminus | Core | C-terminus | 44 |
| $Y_{1d}$ | | Whole Structure | – | – | 0 |
| $Y_{1e}$ | E | Termini | Core | – | 24 |
| $Y_{1f}$ | | N-terminus | Core | C-terminus | 49 |
| $Y_{1h}$ | | Whole Structure | – | – | 0 |
| $Y_{1i}$ | C | Termini | Core | – | 40 |
| $Y_{1j}$ | | N-terminus | Core | C-terminus | 91 |

Table S9: Comparing models with termini of secondary structures separated from the core of the structure only considering secondary structures of 7 codons or longer with the terminus defined as the first and last 3 codons of the structure. H: helix. E: sheet. C: coil.

| Structure Model | Secondary Structure | Groupings I | II | III | ΔDIC |
|---|---|---|---|---|---|
| $Y_{1a}$ | | Whole Structure | – | – | 0 |
| $Y_{1b}$ | H | Termini | Core | – | 7 |
| $Y_{1c}$ | | N-terminus | Core | C-terminus | 33 |
| $Y_{1d}$ | | Whole Structure | – | – | 0 |
| $Y_{1e}$ | E | Termini | Core | – | 45 |
| $Y_{1f}$ | | N-terminus | Core | C-terminus | 88 |
| $Y_{1g}$ | | Whole Structure | – | – | 0 |
| $Y_{1h}$ | C | Termini | Core | – | 49 |
| $Y_{1i}$ | | N-terminus | Core | C-terminus | 103 |

Table S10: Comparing models with termini of secondary structures separated from the core of the structure only considering secondary structures of 8 codons or longer with the terminus defined as the first and last 2 codons of the structure. H: helix. E: sheet. C: coil.

| | | Groupings | | | |
|---|---|---|---|---|---|
| Structure Model | Secondary Structure | I | II | III | ΔDIC |
| $Y_{1a}$ | H | Whole Structure | – | – | 0 |
| $Y_{1b}$ | | Termini | Core | – | 6 |
| $Y_{1c}$ | | N-terminus | Core | C-terminus | 38 |
| $Y_{1d}$ | E | Whole Structure | – | – | 0 |
| $Y_{1e}$ | | Termini | Core | – | 37 |
| $Y_{1f}$ | | N-terminus | Core | C-terminus | 57 |
| $Y_{1g}$ | C | Whole Structure | – | – | 0 |
| $Y_{1h}$ | | Termini | Core | – | 36 |
| $Y_{1i}$ | | N-terminus | Core | C-terminus | 85 |

Table S11: Comparing models with termini of secondary structures separated from the core of the structure only considering secondary structures of 8 codons or longer with the terminus defined as the first and last 3 codons of the structure. H: helix. E: sheet. C: coil.

| | | Groupings | | | |
|---|---|---|---|---|---|
| Structure Model | Secondary Structure | I | II | III | ΔDIC |
| $Y_{1b}$ | H | Termini | Core | – | 0 |
| $Y_{1a}$ | | Whole Structure | – | – | 1 |
| $Y_{1c}$ | | N-terminus | Core | C-terminus | 28 |
| $Y_{1d}$ | E | Whole Structure | – | – | 0 |
| $Y_{1e}$ | | Termini | Core | – | 49 |
| $Y_{1f}$ | | N-terminus | Core | C-terminus | 89 |
| $Y_{1g}$ | C | Whole Structure | – | – | 0 |
| $Y_{1h}$ | | Termini | Core | – | 44 |
| $Y_{1i}$ | | N-terminus | Core | C-terminus | 94 |

Table S12: Comparing models with termini of secondary structures separated from the core of the structure only considering secondary structures of 9 codons or longer with the terminus defined as the first and last 3 codons of the structure. H: helix. E: sheet. C: coil.

| Structure Model | Secondary Structure | Groupings I | II | III | ΔDIC |
|---|---|---|---|---|---|
| $Y_{1b}$ | | Termini | Core | – | 0 |
| $Y_{1a}$ | H | Whole Structure | – | – | 3 |
| $Y_{1c}$ | | N-terminus | Core | C-terminus | 40 |
| $Y_{1d}$ | | Whole Structure | – | – | 0 |
| $Y_{1e}$ | E | Termini | Core | – | 46 |
| $Y_{1f}$ | | N-terminus | Core | C-terminus | 86 |
| $Y_{1g}$ | | Whole Structure | – | – | 0 |
| $Y_{1h}$ | C | Termini | Core | – | 36 |
| $Y_{1i}$ | | N-terminus | Core | C-terminus | 79 |

Table S13: Comparing models with termini of secondary structures separated from the core of the structure only considering secondary structures of 10 codons or longer with the terminus defined as the first and last 3 codons of the structure. H: helix. E: sheet. C: coil.

| Structure Model | Secondary Structure | Groupings I | II | III | ΔDIC |
|---|---|---|---|---|---|
| $Y_{1a}$ | | Whole Structure | – | – | 0 |
| $Y_{1b}$ | H | Termini | Core | – | 13 |
| $Y_{1c}$ | | N-terminus | Core | C-terminus | 63 |
| $Y_{1d}$ | | Whole Structure | – | – | 0 |
| $Y_{1e}$ | E | Termini | Core | – | 43 |
| $Y_{1f}$ | | N-terminus | Core | C-terminus | 72 |
| $Y_{1g}$ | | Whole Structure | – | – | 0 |
| $Y_{1h}$ | C | Termini | Core | – | 33 |
| $Y_{1i}$ | | N-terminus | Core | C-terminus | 74 |

Table S14: Effects of minimum $\alpha$-helix (DSSP H) length and termini length for evaluating differences in selection at termini vs. the core of the structure. $3_10$-helices (DSSP G) and $\pi$-helices (DSSP I) are excluded.

| Minimum $\alpha$-helix Length | Terminus Length | Category 1 | Category 2 | $\Delta$DIC |
|---|---|---|---|---|
| 4 | 2 | Core | Termini | 0 |
| | | Whole Structure | – | 6 |
| 5 | 2 | Core | Termini | 0 |
| | | Whole Structure | – | 2 |
| 6 | 2 | Whole Structure | | 0 |
| | | Core | Termini | 3 |
| 7 | 2 | Whole Structure | – | 0 |
| | | Core | Termini | 21 |
| 8 | 2 | Whole Structure | – | 0 |
| | | Core | Termini | 14 |
| 9 | 2 | Whole Structure | – | 0 |
| | | Core | Termini | 13 |

Table S15: Comparing models for *E. coli* with termini of secondary structures separated from the core of the structure only considering secondary structures of 4 codons or longer with the terminus defined as the first and last 2 codons. H: helix. E: sheet. C: coil.

| | | Groupings | | | |
|---|---|---|---|---|---|
| Structure Model | Secondary Structure | I | II | III | $\Delta$DIC |
| $E_{1b}$ | | Termini | Core | – | 0 |
| $E_{1c}$ | H | N-terminus | Core | C-terminus | 22 |
| $E_{1a}$ | | Whole Structure | – | – | 91 |
| $E_{1f}$ | | N-terminus | Core | C-terminus | 0 |
| $E_{1d}$ | E | Whole Structure | – | – | 48 |
| $E_{1e}$ | | Termini | Core | – | 88 |
| $E_{1g}$ | | Whole Structure | – | – | 0 |
| $E_{1h}$ | C | Termini | Core | – | 3 |
| $E_{1i}$ | | N-terminus | Core | C-terminus | 8 |

Table S16: Comparing models for *E. coli* with termini of secondary structures separated from the core of the structure only considering secondary structures of 5 codons or longer with the terminus defined as the first and last 2 codons. H: helix. E: sheet. C: coil.

| Structure Model | Secondary Structure | Groupings I | II | III | ΔDIC |
|---|---|---|---|---|---|
| $E_{1b}$ | H | Termini | Core | – | 0 |
| $E_{1c}$ | | N-terminus | Core | C-terminus | 15 |
| $E_{1a}$ | | Whole Structure | – | – | 75 |
| $E_{1f}$ | E | N-terminus | Core | C-terminus | 0 |
| $E_{1d}$ | | Whole Structure | – | – | 24 |
| $E_{1e}$ | | Termini | Core | – | 61 |
| $E_{1g}$ | C | Whole Structure | – | – | 0 |
| $E_{1i}$ | | N-terminus | Core | C-terminus | 5 |
| $E_{1h}$ | | Termini | Core | – | 8 |

Table S17: Comparing models for *E. coli* with termini of secondary structures separated from the core of the structure only considering secondary structures of 7 codons or longer with the terminus defined as the first and last 2 codons. H: helix. E: sheet. C: coil.

| Structure Model | Secondary Structure | Groupings I | II | III | ΔDIC |
|---|---|---|---|---|---|
| $E_{1b}$ | H | Termini | Core | – | 0 |
| $E_{1c}$ | | N-terminus | Core | C-terminus | 14 |
| $E_{1a}$ | | Whole Structure | – | – | 44 |
| $E_{1d}$ | E | Whole Structure | – | – | 0 |
| $E_{1e}$ | | Termini | Core | – | 49 |
| $E_{1f}$ | | N-terminus | Core | C-terminus | 53 |
| $E_{1g}$ | C | Whole Structure | – | – | 0 |
| $E_{1h}$ | | Termini | Core | – | 18 |
| $E_{1i}$ | | N-terminus | Core | C-terminus | 45 |
